# NRC immune receptor networks show diversified hierarchical genetic architecture across plant lineages

**DOI:** 10.1101/2023.10.25.563953

**Authors:** Foong-Jing Goh, Ching-Yi Huang, Lida Derevnina, Chih-Hang Wu

**Author notes:** One-sentence summary: The NRC networks show degrees of complexity across asterids, encompassing largely conserved NRC0 networks and diversified family-specific NRC networks.

## Abstract

Plants developed sophisticated immune systems with nucleotide-binding domain and leucine-rich repeat-containing (NLR) proteins to repel invading pathogens. The NRC (NLR required for cell death) family includes helper NLRs that form a complex genetic network with multiple sensor NLRs to provide resistance against pathogens of solanaceous plants. However, the evolution and function of NRC networks outside solanaceous plants is currently unknown. We conducted phylogenomic and macroevolutionary analyses comparing NLRs identified from different asterids lineages and found that NRC networks expanded significantly in most lamiids but not in Ericales and campanulids. Using transient expression assays in *Nicotiana benthamiana*, we show that NRC networks are simple in Ericales and campanulids, but are with high complexity in lamiids. Phylogenetic analyses grouped the NRC helper NLRs into three NRC0 subclades that are conserved, and several family-specific NRC subclades of lamiids that show signatures of diversifying selection. Functional analyses revealed that members of the NRC0 subclades are partially interchangeable, whereas family-specific NRC members in lamiids lack interchangeability. Our findings highlight the distinctive evolutionary patterns of the NRC networks in asterids and provide potential insights into transferring disease resistance across plant lineages.

## Introduction

Plants have evolved intricate immune systems to protect themselves from pathogen invasion. Intracellular nucleotide-binding domain and leucine-rich repeat (NLR) immune receptors play major roles in plant immunity by detecting effector proteins delivered from pathogens (Dodds and Rathjen 2010; Jones et al. 2016; Ngou et al. 2022). NLR activation often results in a form of programmed cell death known as the hypersensitive response, leading to the restriction of pathogen growth. NLRs exhibit a conserved tripartite structure comprised of an N-terminal domain involved in cell death initiation, a central nucleotide-binding domain (NBD) involved in activation, and a C-terminal leucine-rich repeat (LRR) domain involved in ligand binding and NLR self-regulation (Duxbury et al. 2021). NLRs are classified based on their N-terminal domains into TNLs that contain Toll/interleukin-1 receptor/R protein (TIR) domains, RNLs that contain Resistance to Powdery mildew 8 (RPW8)-like coiled-coil (CC) domains, and CNLs that contain G10-type coiled-coil domains (CC_G10_-NLR) or Rx-type coiled-coil domains (CC_Rx_-NLR) (Shao et al. 2016; Duxbury et al. 2021; Kourelis et al. 2021; Lee et al. 2021).

Molecular and genetic studies have categorised NLRs into three groups based on their mode of action: singletons, pairs, and networks (Adachi et al. 2019b; Contreras et al. 2023a). Singleton NLRs can directly or indirectly detect pathogen effectors and initiate downstream immune responses without the assistance of additional NLRs (Contreras et al. 2023s). The CNLs ZAR1 (HopZ-Activated Resistance 1) from *Arabidopsis thaliana* and Sr35 from wheat are two well-studied examples of singleton NLRs (Wang et al. 2015, 2019; Förderer et al. 2022). ZAR1 indirectly recognizes effectors via its RLCK (Receptor-like cytoplasmic kinase) partners and form pentameric membrane-associated resistosome complexes that function as calcium-permeable channels to activate immune responses (Wang et al. 2019; Bi et al. 2021). Sr35 directly binds the stem rust effector AvrS35 through its LRR domain, forming a pentameric resistosome complex similar to ZAR1 (Förderer et al. 2022). NLRs can also function in pairs, in which a sensor NLR, specialised to recognise pathogen effectors, is coupled with a helper NLR that is involved in immune signalling. Paired NLRs often physically interact and exist as linked gene pairs or clusters on the chromosomes, suggesting that they are co-regulated and may have co-evolved exclusively with each other throughout their evolutionary trajectory (Xi et al. 2022). Classical examples of paired NLRs include RRS1/RPS4 of *A. thaliana*, and RGA4/RGA5 and Pik-1/Pik-2 of rice (Ashikawa et al. 2008; Narusaka et al. 2009; Césari et al. 2014; Sohn et al. 2014; Xi et al. 2022). In these pairs, the sensor NLRs often contain integrated domains (ID) that play critical roles in sensing pathogen effectors; for example, the WRKY domain in RRS1 and HMA (Heavy-Metal-Associated) domain in RGA5 and Pik-1 (Cesari et al. 2013; Maqbool et al. 2015; Sarris et al. 2015; Marchal et al. 2022).

NLRs can also function in networks, in which multiple sensor NLRs that detect different pathogen effectors signal through a set of helper NLRs to mediate immunity (Wu et al. 2017, 2018). NRG1 (N-Required Gene 1) and ADR1 (Activated Disease Resistance 1), two RNL-type helper NLRs, are required by multiple sensor NLRs for immune signalling (Castel et al. 2019; Saile et al. 2020). In Arabidopsis, the TNLs RPS4/RRS1, RPP2, and RPS6, preferentially signal through NRG1, whereas the CNLs RPS2 and RPP4 preferentially signal through ADR1, supporting the idea that the two groups of helper NLRs show complex genetic redundancy (Saile et al. 2020). Similar to NRG1 and ADR1, NRCs (NLR-required for cell death) are required by multiple Solanaceae sensor NLRs. Within the NRC network, the NRC-dependent sensor NLRs detect different pathogen effectors and require partially redundant helper NRCs for immune response. For example, sensor NLRs Rpi-blb2, Mi-1.2, and R1 transmit signals via NRC4 to trigger cell death. The sensor NLR Prf and Rpi-amr1 operates with either NRC2 or NRC3, while Sw5b, R8, Rx, Bs2, and Rpi-amr3 can signal redundantly through NRC2, NRC3, or NRC4 (Wu et al. 2016, 2017; Chen et al. 2021; Witek et al. 2021; Wu and Kamoun 2021; Lin et al. 2022; Zhang et al. 2024).

Phylogenetically, NRCs and NRC-dependent sensor NLRs form the NRC superclade, which is a subgroup of CC_Rx_-NLR (Wu et al. 2017; Kourelis et al. 2021; Contreras et al. 2023a). The NRC superclade is present across the asterids and in some Caryophyllales, but is not present in monocots or rosids. This suggests that the ancestral sequences of the NRC superclade arose before the diversification of Caryophyllales and asterids (Wu et al. 2017). A recent report defined NRC0, an NRC helper NLR, as the only conserved family member found in various asterids (Sakai et al. 2023). NRC0 often exists in a gene cluster with NRC0-dependent sensor NLRs, strengthening the hypothesis that the NRC superclade originated from an ancient helper-sensor NLR gene cluster (Wu et al. 2017; Sakai et al. 2023). Another intriguing evolutionary aspect of the NRC superclade is the presence of noncanonical extended N-terminal domains (exNT), which exist before the CC domain in some clades of sensor NLRs (Seong et al. 2020). Although the detailed evolutionary history of the sensor NLRs is not clear, the exNT were also found in sugar beet, suggesting that the exNT of NRC-dependent sensor NLRs emerged between the common ancestor of asterids and Amaranthaceae (Caryophyllales) (Seong et al. 2020).

Upon sensor NLR activation, NRCs form higher-molecular-weight complexes that localize to the plasma membrane and likely act as calcium-permeable channels similar to ZAR1 (Duggan et al. 2021; Ahn et al. 2023; Contreras et al. 2023b). The first alpha helixes of most NRCs possess a conserved α1 helix domain known as the MADA motif that plays a major role in executing cell death (Adachi et al. 2019a). The MADA motif is conserved in ZAR1 and many other singleton NLRs but is absent in the NRC-dependent sensor NLRs, suggesting that degeneration of the MADA motif may be an evolutionary feature associated with the functional diversification of the NRC superclade members (Adachi et al. 2019a). Interestingly, members of the NRC family have also evolved functions beyond triggering cell death induced by sensor NLRs. The cell surface receptor Cf-4 induces NRC3-dependent hypersensitive responses upon detection of the plant pathogen effector AVR4, indicating that the NRC superclade not only functions in intracellular receptor defence but also contributes to cell surface receptor-mediated defences (Kourelis et al. 2022). Furthermore, NRCx, an unusual NRC family member lacking a functional MADA motif, modulates NRC2/NRC3-mediated cell death in *Nicotiana benthamiana* plants. NRCx functions as a negative regulator of cell death execution, thus, playing a key role in maintaining homeostasis within the NRC network (Adachi et al. 2023a).

NLRs represent one of the most diverse protein families in angiosperms, with many species encoding large and diverse repertoires of NLR genes in the genome (Barragan and Weigel 2021). As they play key roles in the survival of a plant species under emerging pathogen pressures, *NLR* genes are known to show distinguished signs of rapid evolution, even within a single species (Kuang et al. 2004; Jacob et al. 2013; Van De Weyer et al. 2019). Given that NLRs exhibit a high turnover rate, the birth-and-death model has been proposed to describe the evolutionary process of *NLR* genes (Michelmore and Meyers 1998). In this model, the emergence of new NLRs occurs through repetitive cycles of gene duplication. Some genes are maintained in the genome and develop novel capabilities to detect pathogens, while others are either lost or nonfunctionalized due to the accumulation of deleterious mutations (Michelmore and Meyers 1998; Barragan and Weigel 2021). These dynamic evolutionary patterns allow the plant immune system to effectively adapt to the rapidly evolving effector repertoires of pathogenic microbes. Indeed, some NLRs have been lost in certain plant lineages but highly expanded in other plant lineages (Shao et al. 2016; Martin et al. 2023). TNLs and NRG1, along with their associated signalling partner SAG101, have been lost in monocots, some dicots, and some Magnoliids (Tarr and Alexander 2009; Collier et al. 2011; Shao et al. 2016; Liu et al. 2021; Wu et al. 2021). However, TNL and RNL are more highly expanded and diversified in Gymnosperms and Rosids (Terefe-Ayana et al. 2012; Shao et al. 2016; Van De Weyer et al. 2019; Van Ghelder et al. 2019; Woudstra et al. 2023). CNLs are in general more abundant in the genomes of most angiosperms, whereas TNLs have been lost frequently in dicots (Liu et al. 2021). Studies of wild tomato and Arabidopsis demonstrated a wide range of NLR polymorphisms within single-species populations, suggesting a correlation between plant adaptation to pathogens and NLR diversity (Stam et al. 2019; Van De Weyer et al. 2019). These NLR polymorphisms may contribute to species-specific signalling pathways in NLR-mediated resistance. For example, CaRpi-blb2, a pepper homolog of wild tomato Rpi-blb2, initiates cell death via NRC8 and NRC9 which were identified only in pepper but not other solanaceous species, highlighting the contribution of lineage-specific NLR clades that emerged recently in resistance against pathogens (Oh et al. 2023).

The availability and improvement of genome databases have enabled the analyses of NLR genes from many model and non-model species, leading to a more comprehensive understanding of NLR evolution and diversity (Thind et al. 2018; Van De Weyer et al. 2019; Barragan and Weigel 2021; Kourelis et al. 2021; Ence et al. 2022). Plant NLRs likely originated from the common ancestor of green algae before rapidly diversifying and evolving in land plants (Gao et al. 2018; Ortiz and Dodds 2018; Shao et al. 2019). The discovery of the MAEPL motif and the MADA motif, which are both crucial in executing cell death, provides evidence of the shared function of the CC domain between non-flowering and flowering plants, strengthening the concept of the origin of NLR from common green algae (Adachi et al. 2019a; Chia et al. 2022). While a high number of NLR genes could offer potential survival advantages, several factors have been proposed to influence the number of NLR genes in a particular plant species. Adaptations to aquatic, parasitic, and carnivorous lifestyles correlated to low NLR numbers among closely related species (Baggs et al. 2020; Liu et al. 2021). For example, the aquatic plant duckweed (*Lemna minor*) has only 11 NLR genes, and the carnivorous aquatic plant *Utricularia gibba* has completely lost all NLR genes (Baggs et al. 2020; Liu et al. 2021). To date, close to 500 NLRs from 31 genera belonging to 11 orders of flowering plants have been experimentally validated (Kourelis et al. 2021). These NLRs emerged in different plant species and likely diversified to function against various pathogens. The diversity of NLRs found in plants may reflect the outcome of their long-standing coevolutionary arms race with pathogens (Upson et al. 2018).

Asterids constitute a highly diverse group of angiosperms and encompass numerous economically important crops. The NRC superclade exists in several asterids species, suggesting that NRC networks may play roles in disease resistance in these plants (Wu et al. 2017; Sakai et al. 2023). Understanding the diversity of NRC networks across asterids may provide useful information on lineage-or species-specific sensor-helper NLR connections, which could be useful for disease resistance breeding and help facilitate interspecies resistance gene transfer. However, our knowledge of the evolutionary history and diversity of NRC networks beyond Solanaceae is limited. To address this, we performed macroevolutionary analyses of NRC networks across asterids and experimentally validated the sensor-helper dependency using heterologous expression in *N. benthamiana*. We found that the NRC superclade displays distinct duplication and expansion patterns in the three asterids lineages (Ericales, campanulids, and lamiids). Combined with the results from transient expression assays in *N. benthamiana*, we revealed that the NRC networks in Ericales, campanulids, and lamiids show different complexity and hierarchical structures. The NRC helper NLR family can be further grouped into three NRC0 subclades that are conserved, and several family-specific NRC subclades of lamiids that show signatures of diversifying selection. Further inter-species comparisons revealed that members of the NRC0 subclades were partially interchangeable, as many of them can function with NRC0-dependent sensor NLRs from different lineages. In contrast, members of the family-specific NRC subclades of lamiids lack interchangeability, with only some of the NRCs showing compatibility with sensor NLRs across plant families. Our study sheds light on the unique evolutionary patterns of the NRC networks within the asterids and offers valuable insights into the potential transfer of disease resistance mechanisms across different plants.

## Results

### The NRC superclade is expanded differentially in distinct plant lineages

To understand the evolutionary diversity of the NRC network in plants, we performed comparative phylogenomic analyses of 46 angiosperms, including basal angiosperms, monocots, rosids, Caryophyllales, and asterids (comprising Ericales, campanulids and lamiids) (Supplemental Data Set 1). We used MAST - MEME Suite and 20 previously defined motifs to predict potential NLR encoding sequences from the genome of all these angiosperms (Jupe et al. 2012). After generating the NLR phylogenetic trees of each species using the conserved NB-ARC domain, we grouped these NLRs into four categories including TNL, RNL, CC_G10_-NLR, and the CC_Rx_-NLR. To identify the NRC superclades in these species, we performed additional phylogenetic analyses of the CC_Rx_-NLR using the solanaceous NLRs (NRC, NRC-dependent and NRC-independent NLRs) as references. These second phylogenetic trees classified the CC_Rx_-NLR into the NRC superclade and non-NRC superclade CNLs. Consistent with the previous finding, the NRC superclade was found only in species of some Caryophyllales and most asterids (Supplemental Data Set 2) (Wu et al. 2017).

Next, we calculated the percentage of NRC superclade members relative to the total number of NLRs in these genomes (Fig. 1A). Interestingly, the percentages of NRC superclade members relative to the total number of NLRs show striking variations across different plant lineages. In Caryophyllales, two out of the seven species we analysed contained the NRC superclade (Fig. 1A). Sugar beet (*Beta vulgaris*) and carnation (*Dianthus caryophyllus*) have approximately 4% and 15% of their total NLRs belonging to the NRC superclade, respectively. In Ericales, all three species we analysed contain members of the NRC superclade (Fig. 1A). The percentages of NRC superclade members out of total NLR in kiwifruit (*Actinidia chinensis*), tea (*Camelia sinensis*) and miracle fruit (*Synsepalum dulcificum*) are 7%, 3% and 1%, respectively. In campanulids, the percentage of NLRs belonging to the NRC superclade is variable, with an average of 8% out of total NLRs (Fig. 1A). Among the campanulids analysed, ginseng (*Panax ginseng*) has the lowest percentage of NRC superclade members, with 3% (4 out of 131) of its NLRs belonging to the NRC superclade; candyleaf (*Stevia rebaudiana*) has the highest percentage, with 14% (44 out of 298) of its NLRs belonging to the NRC superclade. In lamiids, the proportions of NRC superclade members among total NLRs are highly variable (Fig. 1A). Overall, the number of NRC superclade members in most lamiids is higher than that of Caryophyllales, Ericales, and campanulids. In most lamiids species, over 40% of the total NLRs belong to the NRC superclade. In solanaceous plants (such as tomato *Solanum lycopersicum* and *N. benthamiana*), where the NRC superclade and NRC network were originally described (Wu et al., 2017), around 50% of total NLRs belong to the NRC superclade. In contrast, over 75% of total NLRs from the three related *Ipomoea* species (sweet potato, *Ipomoea batatas*; threefork morning glory, *Ipomoea trifida*; and littlebell, *Ipomoea triloba*) belong to the NRC superclade (Fig. 1A). Remarkably, Asiatic witchweed (*Striga asiatica*), a parasitic plant, has the highest percentage (89%) of NRC superclade members among all analysed plant species, while dodder (*Cuscuta australis*), another parasitic plant, has the lowest percentage (14%) of NRC superclade members among lamiids, with only one out of seven NLRs belonging to the superclade. Hardy rubber tree (*Eucommia ulmoides*) was the only lamiids species where the NRC superclade was not found (Fig. 1A). These results suggest that the NRC superclade shows very different evolutionary trajectories among plant lineages.

**Figure 1.**
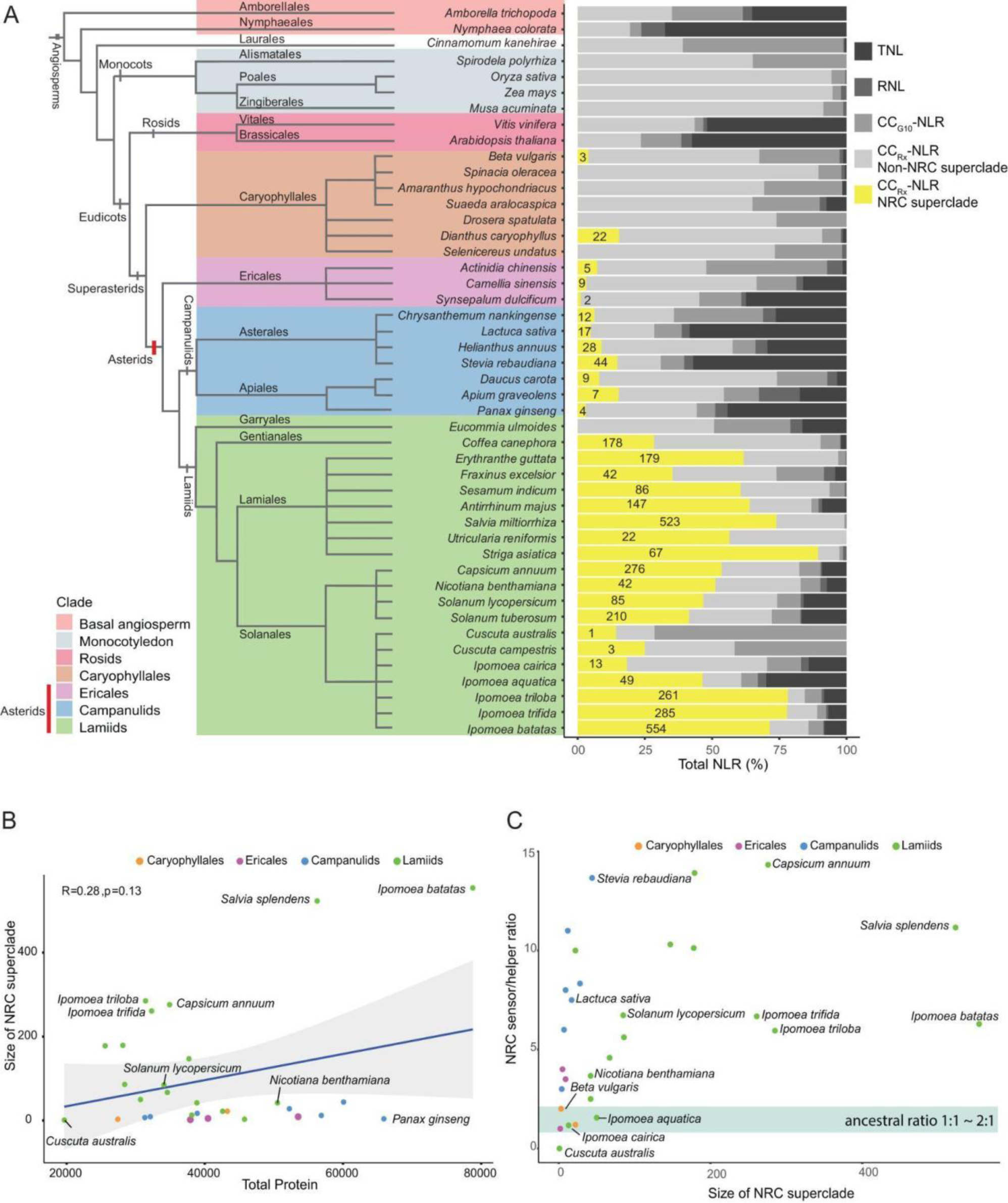
The expansion of the NRC superclade is a unique feature of lamiids. **A)** (Left) Phylogenetic tree of angiosperm species, modified from The Angiosperm Phylogeny Group (2016). (Right) Percentage and number of different types of NLRs identified from each species. Numbers in the stacked bar chart indicate the sizes of NRC superclade members in the plant species. **B)** Correlation between the size of the NRC superclade and the number of total protein-coding genes. **C)** Comparisons of the NRC sensor-helper ratio and the size of the NRC superclade. The highlighted area indicates the ancestral ratio of the NRC sensor and helper, which is close to 1:1 to 2:1. Plant species from different lineages are labelled with different colours.

To understand the degree to which the expansion of the NRC superclade and other NLRs correlates with genome expansion, we compared the size of the genome or the number of protein-coding genes with the number of NLRs or the size of the NRC superclade of each species. The data revealed that there was no correlation between genome size with the amounts of NLRs or the size of the NRC superclade (Supplemental Fig. S1A and S1B), but there was a weak correlation between NLRs and the numbers of total protein-coding genes (R=0.36, p=0.052) (Supplemental Fig. S1C). Additionally, the size of the NRC superclade was very weakly correlated with total protein-coding genes (R=0.28, p=0.13) (Fig. 1B), suggesting that the overall evolutionary pattern of total NLRs is not well aligned with that of the NRC superclade in these species. To further compare the features of the NRC superclade and NLR expansion in different plant lineages, we assessed the number of total NLR/total protein-coding genes and the size of the NRC superclade/total protein-coding genes. Most lamiids had higher ratios of total NRCs to total protein-coding genes compared to other plant lineages (Supplemental Fig. S1D). Based on these analyses, we concluded that the expansion of the NRC superclade is a unique characteristic of lamiids.

The NRC superclade encompasses two types of NLRs, namely the NRC-dependent sensor NLRs (NRC-S) and the NRC helper NLRs (NRC-H). Since the NRC superclade originated from a linked NLR pair or cluster, we expect the ancestral sensor-helper ratio of the NRC superclade to be around 1:1 or 2:1. To gain further insights into whether the sensor and helper NLRs within the NRC superclade have expanded differentially during evolution, we calculated the NRC sensor-helper ratio for all NRC superclade containing plant species (Fig. 1C). Most of the species have NRC sensor-helper ratios greater than 2:1, suggesting that the NRC-S experienced more frequent duplication events in comparison to NRC-H. In Ericales and campanulids, where the NRC superclade constitutes only a small portion of total NLRs, the NRC sensor-helper ratio varied. In Ericales, the NRC sensor-helper ratio ranged from 1:1 to 4:1, while in campanulids this ratio ranged from 3:1 to 14:1. In lamiids, where the NRC superclade is often extensively expanded, most species displayed sensor-helper ratios higher than the ancestral ratio (Fig. 1C). Together, these findings indicate that sensor NLRs are more prone to duplication during evolution compared to helper NLRs, supporting the notion that sensor NLRs require diversification to recognise various pathogens, while helper NLRs are involved in mediating a conserved cell death pathway.

### The NRC network in Ericales is simple

To characterise the NRC superclade in Ericales, we performed phylogenetic analysis on NLRs identified from *A. chinensis*, *C. sinensis*, and *S. dulcificum* using tomato NRC superclade members as a reference (Fig. 2, A and B). The Ericales NRC superclade can be divided into one NRC-H clade and one NRC-S clade. To understand the phylogenetic relationship of the Ericales NRCs with the NRC0 subclade described by Sakai et al. (2023), we performed phylogenetic analyses encompassing all the predicted NRC-H from asterids. We found the Ericales NRC-H clustered together with the NRC0 subclade, but formed its lineage-specific group (Supplemental Fig S2A). Therefore, to differentiate this group of NRCs from the NRC0 subclade, we named the members of this subclade NRC0-Ericales-specific (NRC0-Eri). In several plant species, NRC0 orthologs are physically clustered with the NRC0-dependent sensor NLRs on the chromosome (Sakai et al. 2023). We found a putative NRC-S linked to one of the NRC0-Eri in *C. sinensis* (CsNRC0b-Eri, CSS0008112.1) (Supplemental Fig. S2B and C), but no sensor-helper linkage was found in *A*. *chinensis* or *S. dulcificum*.

**Figure 2.**
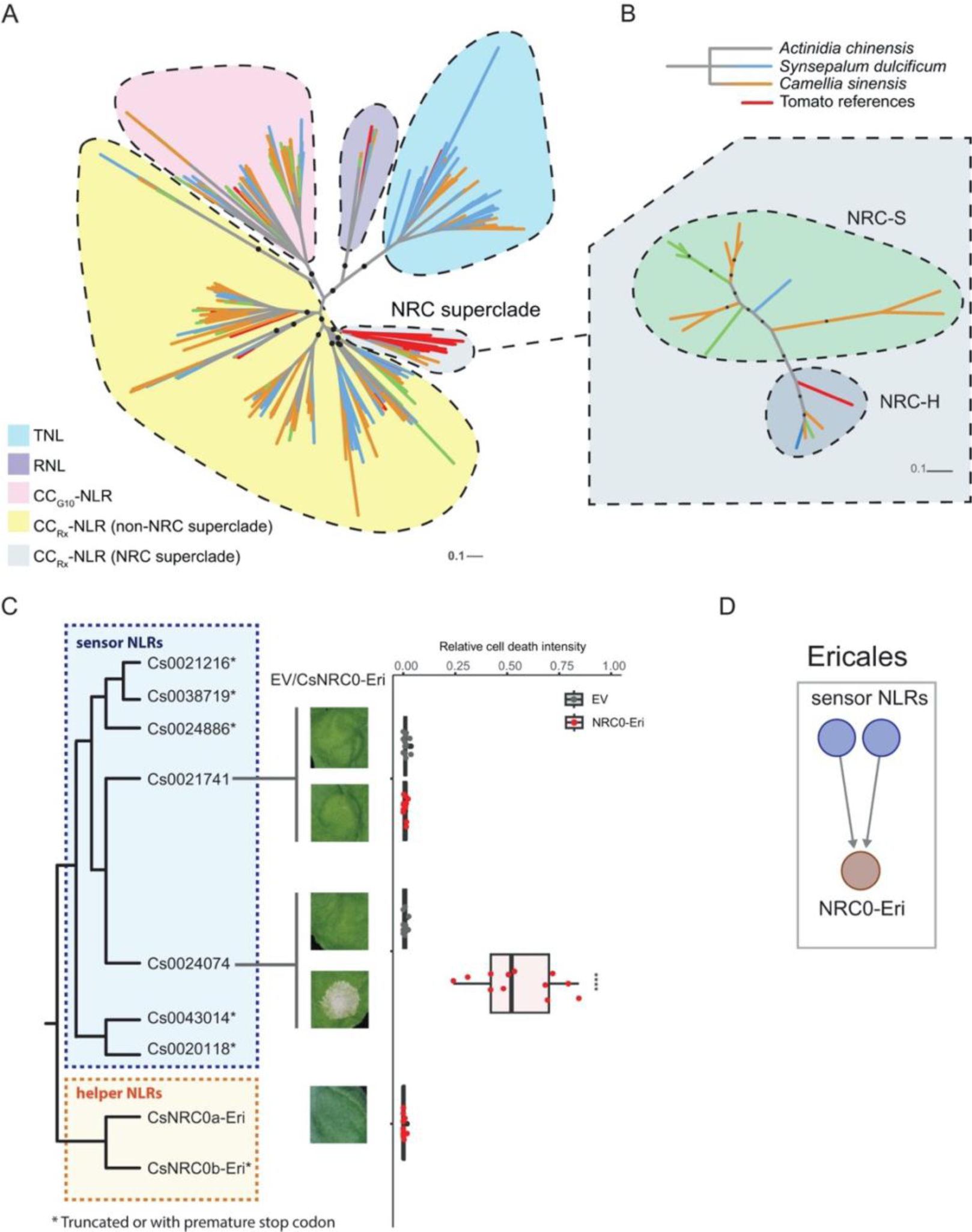
Ericales encode a simple NRC network, with sensor NLRs signalling through NRC0-Eri to initiate cell death. **A)** Phylogenetic analysis of total NLRs from 3 Ericales species, including *A. chinensis*, *S. dulcificum*, and *C. sinensis.* NLRs from different species are indicated with lines of different colours. NLR sequences from the tomato NRC superclade were used as references. **B)** Phylogenetic analysis of the NRC superclade from the 3 Ericales species with tomato NRC0 as reference sequences. For both phylogenetic trees, major branches with bootstrap values higher than 70 are indicated with black dots. The scale bar indicates the evolutionary distance in amino acid substitution per site. **C)** Cell death assay was performed by co-expression of the putative NRC0-Eri-dependent sensor NLR from tea (*C. sinensis*) and tea CsNRC0a-Eri. All sensor NLRs carried the MHD motif (D to V) mutation. The intensity of cell death was analysed at 5 dpi. The dot plot represents the relative cell death intensity based on autofluorescence imaging using UVP ChemStudio PLUS. The bold line in the boxplots represents the medium, the box edges represent the 25th and 75th percentiles, and the whiskers extend to the most extreme data points no more than 1.5x of the interquartile range. Statistical differences were examined by paired Wilcoxon signed rank test (**** = p < 0.0001; * = p < 0.05). **D)** Putative NRC network structure in Ericales where few sensor NLRs signal through Ericales NRC0 (NRC0-Eri) to induce cell death.

To validate that the members of NRC0-Eri function as helper NLRs for the putative sensor NLRs, we cloned NRC0 from *A. chinensis* (AcNRC0-Eri, Achn230571) and *C. sinensis* (CsNRC0-Eri) and performed transient expression assays in *N. benthamiana* together with putative sensor NLRs from the same species. As our cloned CsNRC0b-Eri (CSS0008112.1) contains a premature stop codon in the NB-ARC domain and expression of AcNRC0-Eri alone induced cell death in wild type and *nrc2/3/4*_KO *N. benthamiana,* we focused on CsNRC0a-Eri (CSS0010223.1) for validation of the Ericales NRC network (Supplemental Fig. S2D). Out of the 7 putative NRC-dependent sensor NLRs annotated in the genome of *C. sinensis*, two contained full-length NLR sequence signatures, and the remaining five were truncated, either missing the coiled-coil domains or part of the LRR domains. We cloned the two full-length *C. sinensis* NLRs (Cs0021741 and Cs0024074) and introduced a D to V mutation into their MHD motifs to generate constitutively active variants (Cs0021741^DV^ and Cs0024074^DV^). Co-expression of Cs0024074^DV^, but not Cs0021741^DV^, with CsNRC0a-Eri induced cell death in *N. benthamiana.* Cell death was not observed when either Cs0024074^DV^ or CsNRC0a-Eri were expressed alone, indicating that CsNRC0a-Eri is required for Cs0024074 cell death activity (Fig. 2C). Overall, our data suggests that CsNRC0a-Eri functions as a helper NLR. Furthermore, the NRC network of Ericales is simple, with few sensor NLRs signalling through one or two NRC0 homologs (Fig. 2D).

### NRCs duplicated in some campanulids, forming an NLR network with partially redundant nodes

To characterise the NRC superclade in campanulids, we performed phylogenetic analyses of NLRs identified from seven species, including *Chrysanthemum nankingense*, lettuce (*Lactuca sativa*), sunflower (*Helianthus annuus*)*, S. rebaudiana,* carrot (*Daucus carota*), celery (*Apium graveolens*), and *P. ginseng*, across two major orders (Asterales and Apiales) (Supplemental Fig. S3A). In line with our previous findings, the campanulids NLRs were classified into several highly supported clades consisting of TNLs, RNLs, CC_G10_-NLRs and CC_Rx_-NLRs, which include the NRC superclade (Fig. 3A and Supplemental Fig. S3A). Phylogenetic analysis revealed that the campanulids NRC superclade is divided into multiple clades, including one NRC-H clade and several putative NRC-S clades (Fig. 3B). The NRC-H clade can be further divided into the NRC0 subclade, which contains sequences from both Asterales, Apiales and the reference tomato NRC0, as well as another subclade which contains sequences only from the Asterales (Fig. 3B). Based on the phylogenetic tree of NRC-H from asterids, this subclade clusters together with NRC0-Eri and the NRC0 subclade defined by Sakai et al. (2023) (Supplemental Fig. S2A). Therefore, we named this group the NRC0-Asterales-specific (NRC0-Ast) subclade.

**Figure 3.**
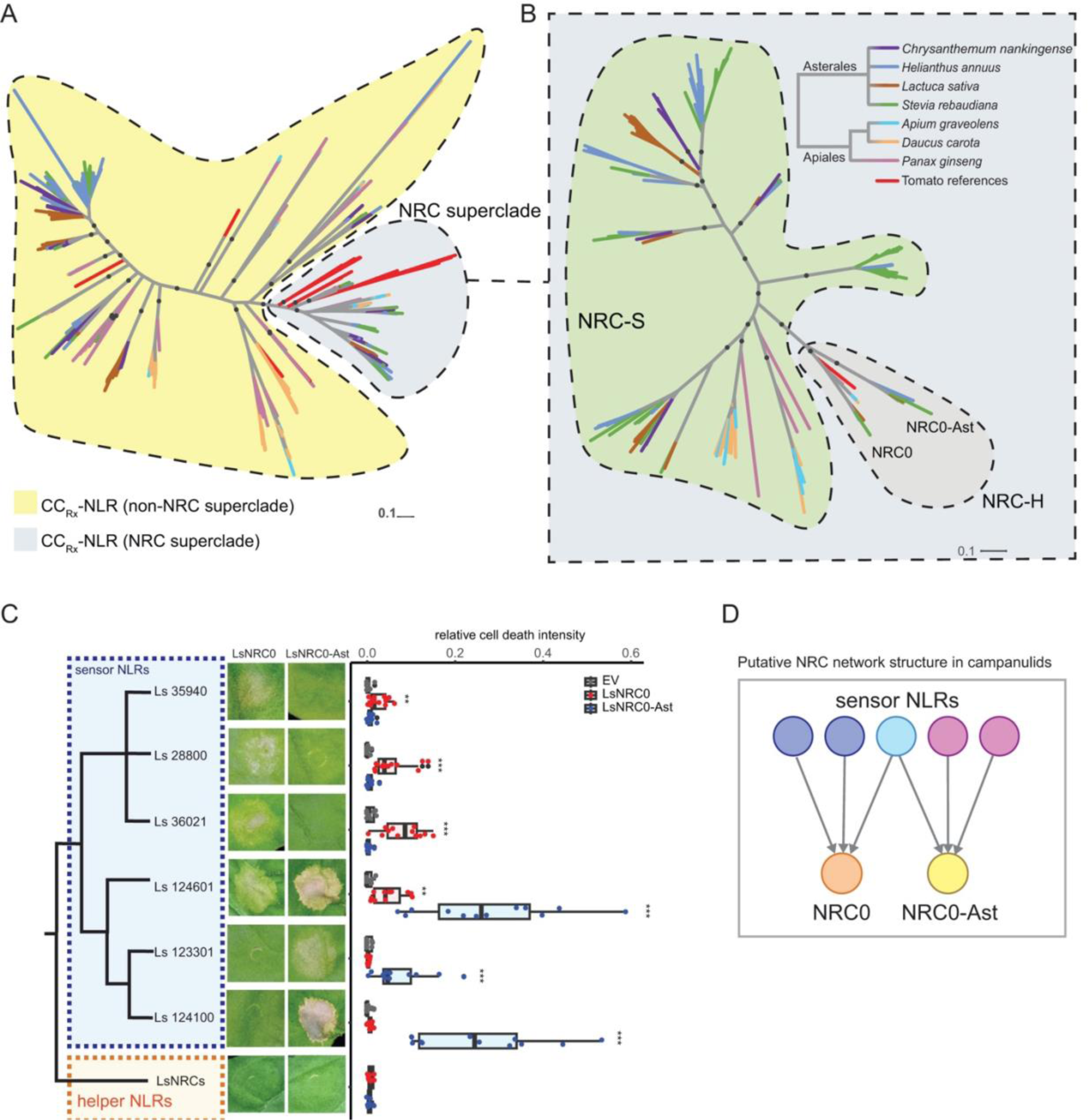
Campanulids show an NRC network with two partially redundant NRC nodes. **A)** Phylogenetic analysis of CCRx-CNLs from 7 species of campanulids, including *C. nankingense*, *H. annuus*, *L. sativa*, *S. rebaudiana*, *A. graveolens*, *D. carota*, and *P. ginseng*. **B)** Phylogenetic analysis of the NRC superclade from 7 species of campanulids using tomato NRC0 as references. Major branches with bootstrap values > 70 are indicated with black dots in both (A) and (B). The scale bar indicates the evolutionary distance in amino acid substitution per site. **C)** Cell death assay results of NRC-dependent sensor NLRs co-expressed with the lettuce NRCs in *N. benthamiana* at 5 dpi. All sensor NLRs carried the MHD motif (D to V) mutation. As controls, helper NLR LsNRCs (LsNRC0 or LsNRC0-Ast) were infiltrated without sensor NLRs. The dot plot represents the cell death quantification analysed by UVP ChemStudio PLUS. The bold line in the boxplots represents the medium, the box edges represent the 25th and 75th percentiles, and the whiskers extend to the most extreme data points no more than 1.5x of the interquartile range. Statistical differences were examined by paired Wilcoxon signed rank test (* = p < 0.05, ** = p < 0.01, *** = p < 0.001, **** = p < 0.0001). **D)** Putative NRC network structure in campanulids where some NRC-S signal through NRC0, some NRC-S signal through NRC0-Ast, and some signal through both NRCs to induce HR.

To determine the genetic structure of the campanulids NRC network, we selected *D. carota* from Apiales and *L. sativa* from Asterales as representative species. The carrot NRC superclade is comprised of DcNRC0 (DCAR_023561) and eight sensor NLRs falling into two subgroups (Fig. 3B and Supplemental Fig. S3B). Out of the eight sensor NLRs, two of them (Dc23557 and Dc23650) are located in a gene cluster together with DcNRC0 (Supplemental Fig. S3, C and D). We cloned four putative sensor NLRs, including Dc23557 and Dc23650, and introduced a D to V mutation into their MHD motifs and then co-infiltrated them independently with DcNRC0. While two of the sensor NLRs (Dc11400^DV^ and Dc12621^DV^) did not induce cell death with DcNRC0, Dc23650^DV^ and Dc23557^DV^ induced different intensities of cell death in the presence of DcNRC0 (Supplemental Fig. S3E). The lettuce NRC superclade is composed of LsNRC0 (Lsat_1_v5_gn_3_132141.1), LsNRC0-Ast (Lsat_1_v5_gn_8_124621), and 15 sensor NLRs. Most of these sensor NLRs are clustered into two sub-groups (Supplemental Fig. S4A). Interestingly, LsNRC0-Ast is located in proximity to some of the sensor NLRs whereas LsNRC0 is located on a different chromosome (Supplemental Fig. S4, B and C). Similar to the previous experiment, we introduced autoactive mutations (D to V) into the MHD motif of the sensor NLRs, and then performed cell death assays by co-agroinfiltration with either LsNRC0 or LsNRC0-Ast on *N. benthamiana* leaves. Three members of the sensor NLRs within the same sub-group (Ls35940^DV^, Ls28800^DV^, and Ls36021^DV^) induced cell death when co-expressed with LsNRC0 but not LsNRC0-Ast. Two putative sensor NLRs within another sub-group (Ls123301^DV^ and Ls124100^DV^) induced cell death when co-expressed with LsNRC0-Ast but not LsNRC0. (Fig. 3C). Interestingly, one of these putative sensor NLRs (Ls124601^DV^) induced cell death when coexpressed with either LsNRC0 or LsNRC0-Ast (Fig. 3C). These results indicate that the sensor NLRs of lettuce are divided into the NRC0-dependent and NRC0-Ast-dependent groups, with some sensor NLRs capable of signalling through both NRCs (Fig. 3, C and D). Altogether, these results suggest that, in campanulids, some orders such as Apiales (carrot) harbour a simple NRC network composed of NRC0 and matching sensor NLRs, whereas the NRC network in other orders such as Asterales (lettuce) contain at least two partially redundant NRCs, namely NRC0 and NRC0-Ast, and the matching sensor NLRs (Fig. 3D).

To gain insight into the interchangeability of NRC networks across different species of campanulids, we conducted cross-species comparisons using NRC helper and sensor NLR from both carrot and lettuce. We found that two LsNRC0-dependent sensor NLRs (Ls36021^DV^ and Ls124601^DV^) trigger cell death when co-expressed with DcNRC0 (Supplemental Fig. S5, A and B). The sensor NLR Ls28800^DV^ triggered weak but statistically significant cell death when co-expressed with DcNRC0 (Supplemental Fig. S5, A and B). The lettuce sensor NLRs Ls124041^DV^ and Ls123301^DV^, which triggered cell death when co-expressed with LsNRC0-Ast, did not trigger cell death when co-expressed with DcNRC0 (Supplemental Fig. S5, A and B). Interestingly, carrot sensor NLR Dc23560^DV^, which induces cell death together with DcNRC0, triggers clear cell death with LsNRC0-Ast and weak but statistically significant cell death with LsNRC0 (Supplemental Fig. S5, C and D). These results revealed that DcNRC0, LsNRC0 and LsNRC0-Ast are partially interchangeable, displaying some compatibility with sensor NLRs across the two species.

### The NRC network is highly expanded in most lamiids

To compare the NRC superclade of different lamiids, we selected 5 lamiids species, including monkeyflower (*Erythranthe guttata*), European ash trees (*Fraxinus excelsior*), coffee (*Coffea canephora*), *S. lycopersicum*, and water spinach (*Ipomoea aquatica*), and generated a phylogenetic tree encompassing their NRC superclade members. This tree showed that, within the NRC superclade, the NRC-H clade was the only well-supported clade that contained NLR sequences from all five species (Fig. 4A). Most of the well-supported NRC-S clades only contained sequences from single species. This is consistent with the view that sensor NLRs are more diversified than helper NLRs in different lamiids lineages. Further phylogenetic analyses revealed that the NRC family is also highly diverse, with only NRC0 from the five selected species forming a well-supported clade (Fig. 4B). The other NRCs across the five species were rarely found to cluster together, indicating that the NRC family has extensively diversified across different lamiids species.

**Figure 4.**
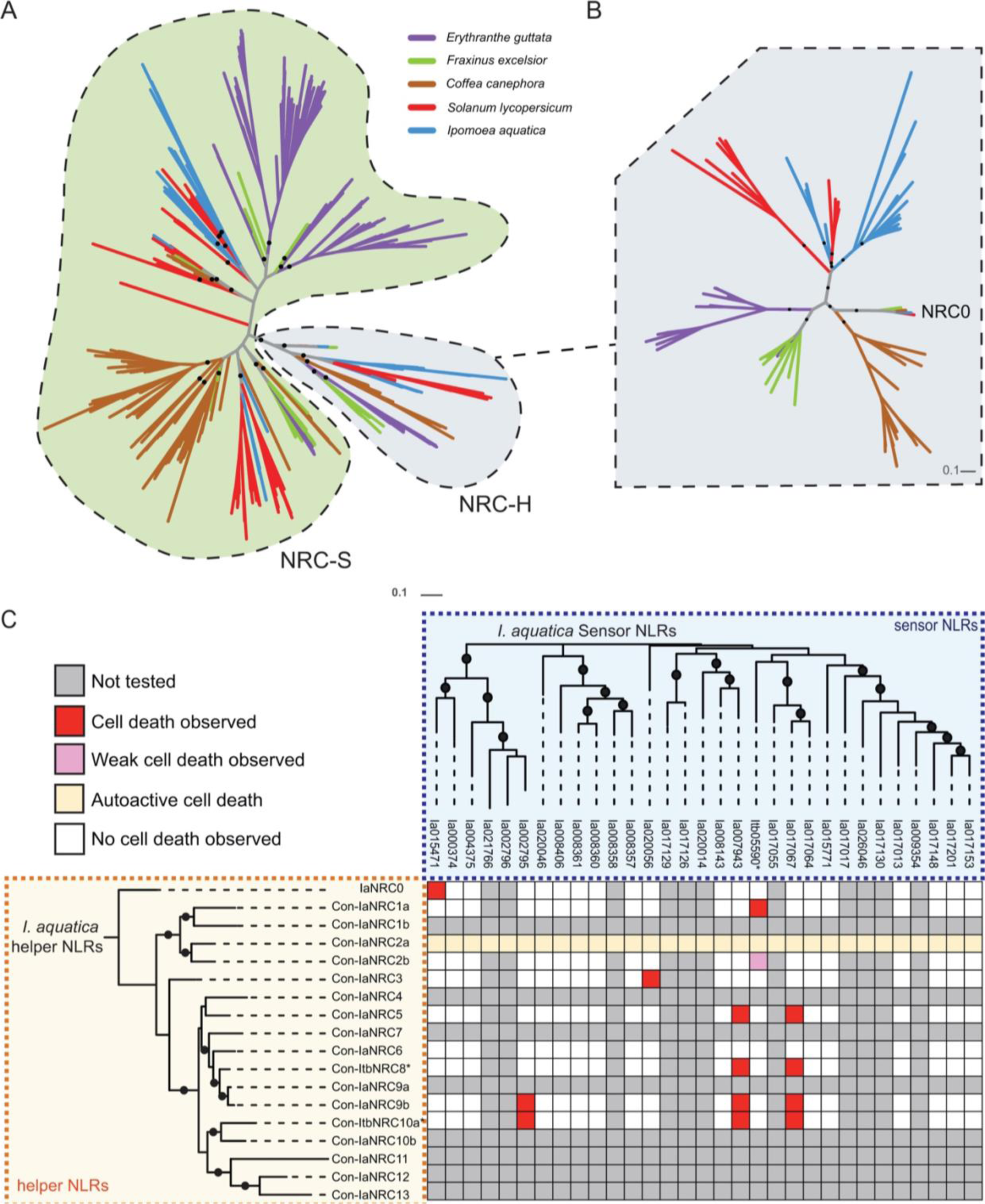
The *Ipomoea* genus possesses a complex NRC network. **A)** Phylogenetic analysis of the NRC superclade of five plant species of lamiids. **B)** Phylogenetic analysis of the NRC family members across five plant species of lamiids. Major branches with bootstrap values > 70 are indicated with black dots in both (A) and (B). The scale bar indicates the evolutionary distance in amino acid substitution per site. **C)** Cell death matrix of *I. aquatica* putative NRC sensor NLRs co-expressed with NRC helper NLRs. All sensor NLRs carried the MHD motif (D to V) mutation. *Sequences cloned form *I. triloba* instead of *I. aquatica* were used in the analysis. Detailed cell death assay results are provided in Supplemental Fig. S7.

We then sought to reconstitute the NRC network using sequences identified from *Ipomoea* species. The *I. aquatica* NRC superclade consists of 18 NRC-H and 31 NRC-S, which is less expanded than that of *I. triloba*, *I. trifida* and *I. batatas* (Fig. 4C, and Supplemental Data Set 2). Given that many of the orthologs of *I. triloba* NRC-H and NRC-S can be identified from *I. aquatica* in the phylogenetic analysis, we reasoned that *I. aquatica* harbours a simplified NRC network compared to that of *I. triloba* (Supplemental Fig. S6, A and B). Therefore, we used *I. aquatica* (Ia) as our primary species for cloning *Ipomoea* NRC-H/S, and *I. triloba* (Itb) as an alternative species when our attempts at cloning failed. Since most of these NRCs are not orthologous to the previously described solanaceous NRCs, we name these 18 NRCs Convolvulaceae(Con)-Ia/ItbNRCs to differentiate them from NRCs identified from the Solanaceae (Sol). In addition to the IaNRC0 (GWHPABKX015475), some of the Con-IaNRCs are located in gene clusters together with putative NRC-S (Supplemental Fig. S6, C and D). We successfully cloned 10 NRCs and 20 putative NRC-S from *I. aquatica* or *I. triloba* and performed cell death assays as described previously (Supplemental Data Set 3). As expected, the sensor NLR Ia15471^DV^ induced cell death when co-expressed with its physically linked helper NLR, IaNRC0, but not with other Con-IaNRC/ItbNRCs (Fig. 4C and Supplemental Fig. S7). An additional 13 sensor NLR and Con-IaNRC/ItbNRCs pairings were shown to function together, displaying strong or weak cell death (see method for definition) when co-expressed in *N. benthamiana* (Fig. 4C and Supplemental Fig. S7). These results indicate that *I. aquatica* contains two independent NRC networks: one simple network with NRC0, similar to what was observed in Ericales and campanulids; and one complex network with multiple NRCs showing varying degrees of genetic redundancy, similar to the phenomenon observed in Solanaceae (Supplemental Fig. S8).

### Family-specific NRC subclades of lamiids show features of diversifying selection

To gain further insights into the evolution of NRC helper NLRs across asterids, we performed phylogenetic analyses using full-length NRC sequences, with the tomato NRC0-dependent sensor NLR (Sl08230) as an outgroup. This phylogenetic tree classified the NRCs into the NRC0 subclades (NRC0, NRC0-Eri, NRC0-Ast) and several lineage-specific NRC subclades of lamiids, consistent with the previous analysis using only the NB-ARC domain (Fig. 5A, Supplemental Fig. S9, and Supplemental Fig. S2). In lamiids, the NRC subclades grouped together in a manner largely consistent with their respective family taxonomy (Fig. 5A, Supplemental Fig. S9, and Supplemental Fig. S2). Consequently, we referred to these clusters as family-specific NRC subclades. Next, we performed an adaptive branch-site REL test for episodic diversification (aBSREL), and found signatures of diversifying selection in several of the lamiids family-specific NRC subclades, but not in the NRC0 subclades (Fig. 5A, Supplemental Fig. S9, and Supplemental Data Set 4). Furthermore, NRC0 subclades showed shorter branch lengths in general, whereas the family-specific NRC subclades showed more variable and longer branch lengths (Fig. 5, B and C, and Supplemental Fig. S9). These results suggest that the three NRC0 subclades are less diversified than the family-specific NRC subclades of lamiids.

**Figure 5.**
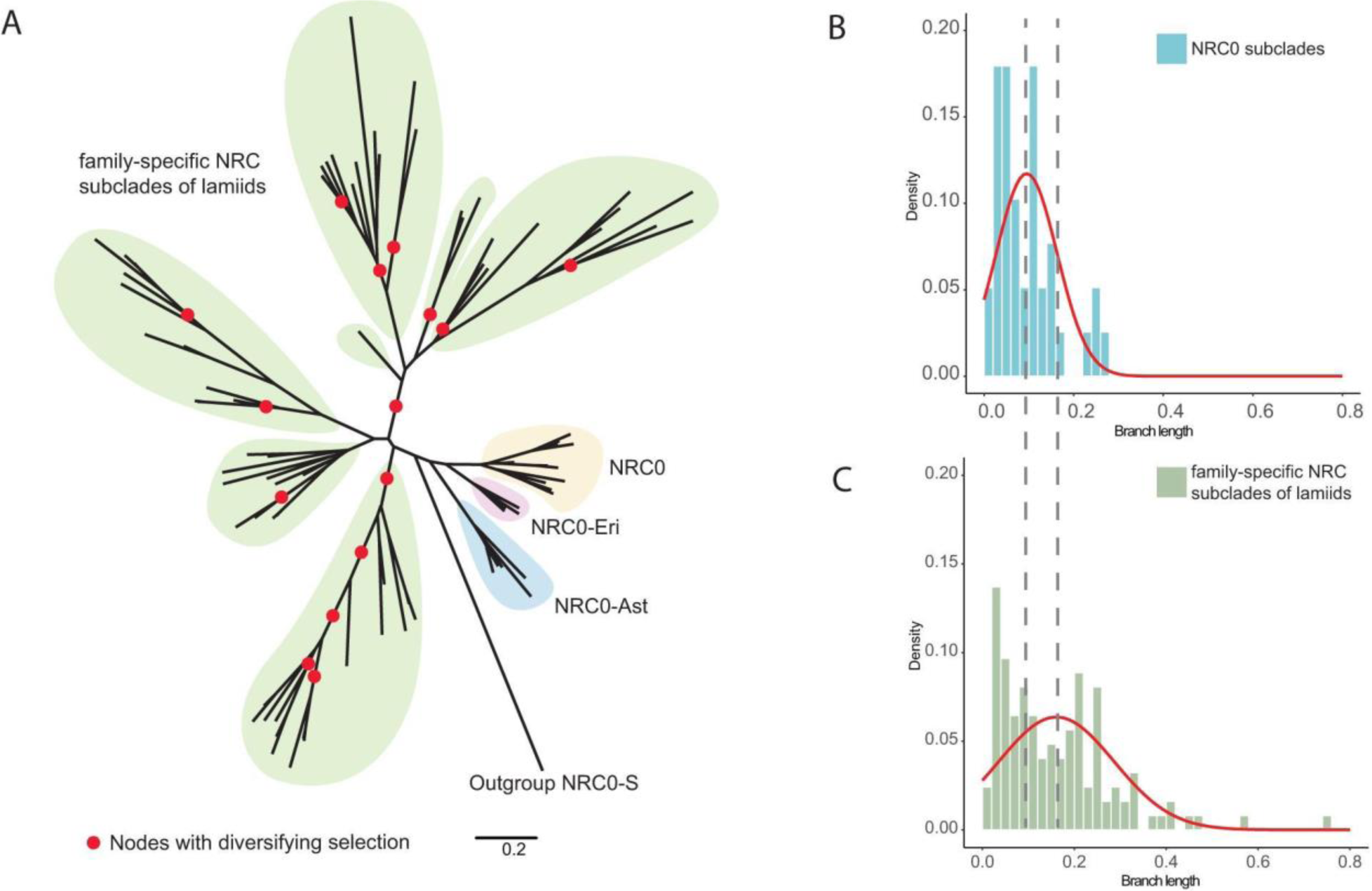
Lamiids family-specific NRC subclades, but not the three NRC0 subclades, show diversifying selection. **A)** Phylogenetic tree based on the full-length sequences of the NRC family from 15 selected asterids species. The aBSREL analysis indicated that 15 out of 81 selected internal branches show episodic diversifying selection. Diversifying selection on branch nodes was assessed using the Likelihood Ratio Test with a significance threshold set at p ≤ 0.05. Detailed phylogeny and results of aBSREL analysis are provided in Supplemental Fig. S9 and Supplemental Data Set 3. The scale bar indicates the evolutionary distance in amino acid substitution per site. **B)** and **C)** Branch length distribution of the three NRC0 subclades and family-specific NRC subclades of lamiids based on the phylogenetic tree in (A). The red lines in both histograms indicate the predicted normal distribution.

### NRC0 subclades across asterids are partially interchangeable

To test the degree to which members of the NRC0 subclades (NRC0, NRC0-Eri, and NRC0-Ast) are interchangeable, we conducted cell death experiments using NRC0 homologs cloned from *C. sinensis, D. carota, L. sativa, S. lycopersicum, I. aquatica*. We found that tea NRC0-dependent sensor NLR Cs24074^DV^ triggered clear cell death when co-expressed with DcNRC0, IaNRC0 and SlNRC0, but no cell death with LsNRC0 or LsNRC0-Ast (Fig. 6, A and E). Out of the two carrot NRC0-dependent sensor NLRs, only one (Dc23650^DV^) induced cell death when co-expressed with LsNRC0-Ast, LsNRC0, IaNRC0, and SlNRC0 (Fig. 6, B and E). The lettuce NRC0-dependent sensor NLRs Ls28800^DV^, Ls35940^DV^ and Ls36021^DV^, which did not induce cell death when expressed with LsNRC0-Ast, induced cell death when co-expressed with most of the NRC0 from the other species, although the cell death with IaNRC0 tends to be weaker for Ls35940^DV^ and Ls36021^DV^ (Fig. 6, C and E). The lettuce NRC0-Ast-dependent sensor NLRs Ls123301^DV^ and Ls124100^DV^ failed to induce cell death with any of the other tested NRC0 variants. Interestingly, the lettuce NRC-S Ls124601^DV^, which can signal through both LsNRC0 and LsNRC0-Ast, is the only NRC-S that activates cell death with all NRC0 variants tested (Fig. 6, C and E). The water spinach sensor NLR Ia15471^DV^ only induced cell death with SlNRC0 or IaNRC0 but not any of the other tested NRC0 variants. Similarly, the tomato NRC0-dependent sensor NLR Sl08230^DV^ only induced cell death with IaNRC0 or SlNRC0 (Fig. 6, D and E). These results indicate that NRC0 homologs exhibit partial interchangeability across different lineages of asterids. Pair-wise sequence comparison of different NRC0 subclades members indicated that LsNRC0-Ast shows lower similarity to other NRC0 subclades members, largely consistent with the results that LsNRC0-Ast displays overall more different sensor compatibility compared to other NRC0 tested (Supplemental Fig. S10A, and Fig. 6E). In the pairwise sensor NLR comparisons, apart from clear homology among some *D. carota* and *L. sativa* sensor NLRs, Ls124601 displayed slightly higher overall similarity to other sensor NLRs included in the analysis (Supplemental Fig. S10B). Intriguingly, Ls124601 was the only sensor NLR that triggered cell death with all the NRC0 homologs tested (Fig. 6C). Together these results suggest that the NRC0 network, across different asterids lineages, is broadly conserved, however, some of the sensor or helper NLRs have diversified and lost their compatibility with each other (Fig. 6E).

**Figure 6.**
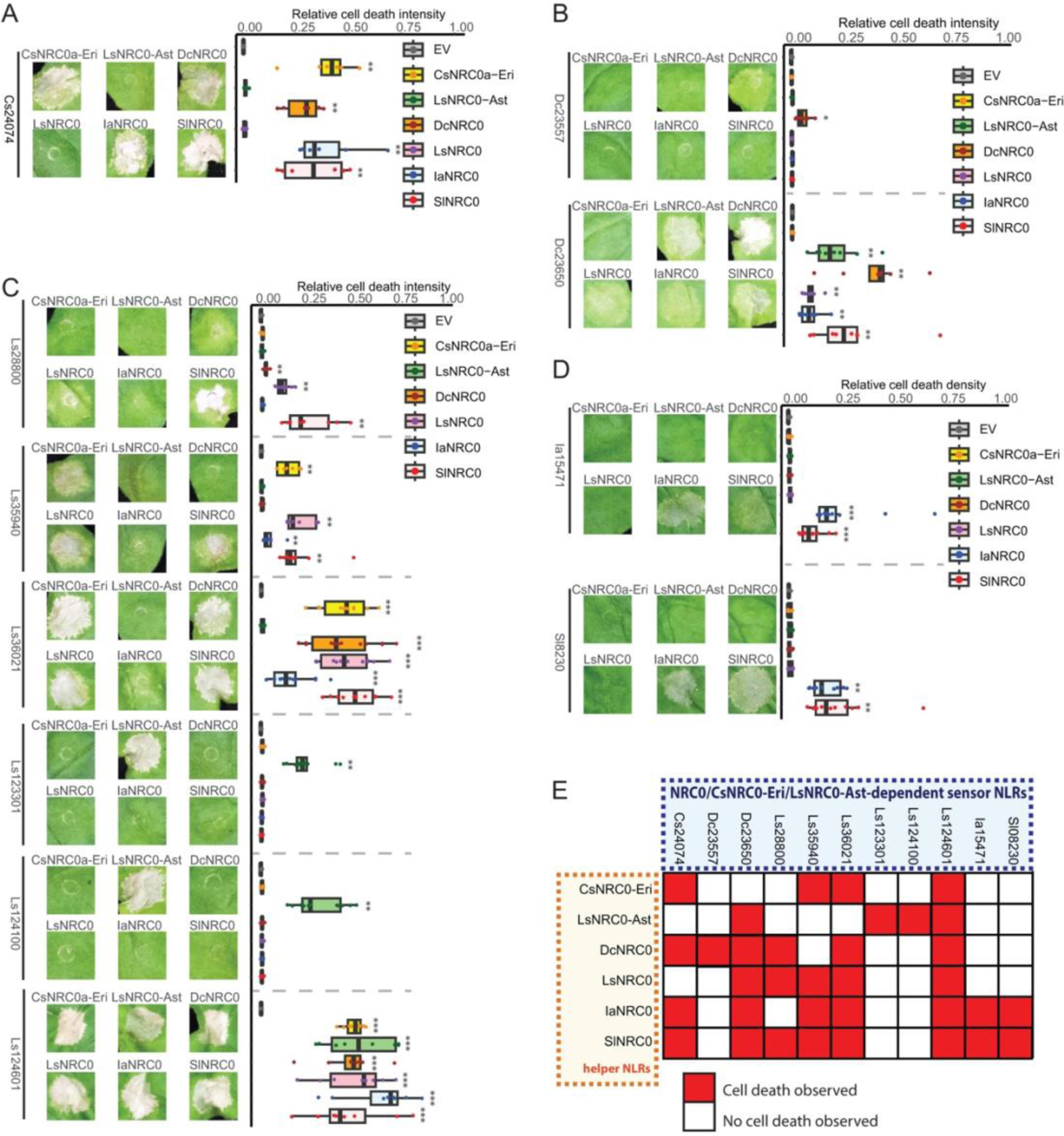
NRC0 subclades from different plant lineages showed varying degrees of interchangeability. **A)** Cell death assay results of tea Cs24074 co-expressed with DcNRC0, LsNRC0, LsNRC0-Ast, IaNRC0, and SlNRC0 in *N. benthamiana* observed at 5 dpi. **(B)** Cell death assay results of carrot sensor NLRs (Dc23557 and Dc23650) or lettuce sensor NLRs (Ls28800, Ls35940, Ls36021, Ls123301, Ls124100, and Ls124601) co-expressed with CsNRC0-Eri, IaNRC0, and SlNRC0 in *N. benthamiana* observed at 5 dpi. **C)** Cell death assay results of water spinach sensor NLR (Ia15471) or tomato sensor NLR (Sl08230) co-expressed with CsNRC0-Eri, DcNRC0, LsNRC0, LsNRC0-Ast, IaNRC0, and SlNRC0 in *N. benthamiana* observed at 5 dpi. All sensor NLRs carried the MHD motif (D to V) mutation. The dot plot represents cell death quantification analysed by UVP ChemStudio PLUS. The bold line in the boxplots represents the medium, the box edges represent the 25th and 75th percentiles, and the whiskers extend to the most extreme data points no more than 1.5x of the interquartile range. Statistical differences were examined by paired Wilcoxon signed rank test. **D)** Matrix of cell death assays of NRC0-dependent sensor NLRs co-expressed with NRC0 homologs from 5 different species of various lineages.

### Sensor and helper NLRs from Solanaceae and Convolvulaceae display some inter-family cross-compatibility

We previously characterised the NRC network of Solanaceae, which consists of conserved NRC homologs across most solanaceous plants (Wu et al. 2017). However, phylogenetic analyses of the NRC superclade here revealed that NRC network members have diversified, and expanded extensively in different families of lamiids (Fig. 4, A and B, and Fig. 5A). This has led us to speculate that sensor and helper NLRs of NRC networks from distinct families in lamiids may share very little or no compatibility. To test this, we co-expressed sensor NLRs from solanaceous plants with NRCs from *Ipomoea* (representing the Convolvulaceae family), and sensor NLRs from *Ipomoea* with NRCs from tomato (representing the Solanaceae family). As expected, most of the solanaceous NLRs, including Bs2, R1, Prf (Pto/AvrPto), and Rpi-blb2, did not trigger cell death when co-expressed with any *Ipomoea* NRCs tested (Supplemental Fig. S11). To our surprise, Rx triggered cell death when co-expressed with Con-IaNRC3, 5, 9, and Con-ItbNRC8 with varying degrees of intensity (Fig. 7, A and B, Supplemental Fig. S12), and Rpi-amr1 triggered cell death when co-expressed with Con-IaNRC9 (Fig. 7, A and B, Supplemental Fig. S12). When we tested the reverse combinations, we found that three of the tested water spinach sensor NLRs failed to induce cell death with the Sol-SlNRCs (Supplemental Fig. S13). In addition to Ia15471^DV^, which we previously showed to signal through Sol-SlNRC0, two *Ipomoea* sensor NLRs (Itb05590^DV^, and Ia17067^DV^) induced cell death with solanaceous Sol-SlNRC1 but not the other Sol-SlNRCs tested (Fig. 7, C and D, Supplemental Fig. S14). However, phylogenetic analysis reveals no clear orthology between Sol-SlNRC1 and any Con-NRCs (Supplemental Fig. S6B), and Itb05590^DV^ and Ia17067^DV^ signalled through different sets of Con-NRCs in the earlier analyses (Fig. 4C). The only group of NRC-S that shows clear homology across these two species are the NRC0-dependent sensor NLRs (Supplemental Fig. S15). This suggests that, despite having family-specific NRC networks, some cross-compatibility between sensor and helper NLRs exists between Solanaceae and Convolvulaceae. Further studies are required to provide mechanistic explanations of this NRC-S and NRC-H inter-family cross-compatibility.

**Figure 7.**
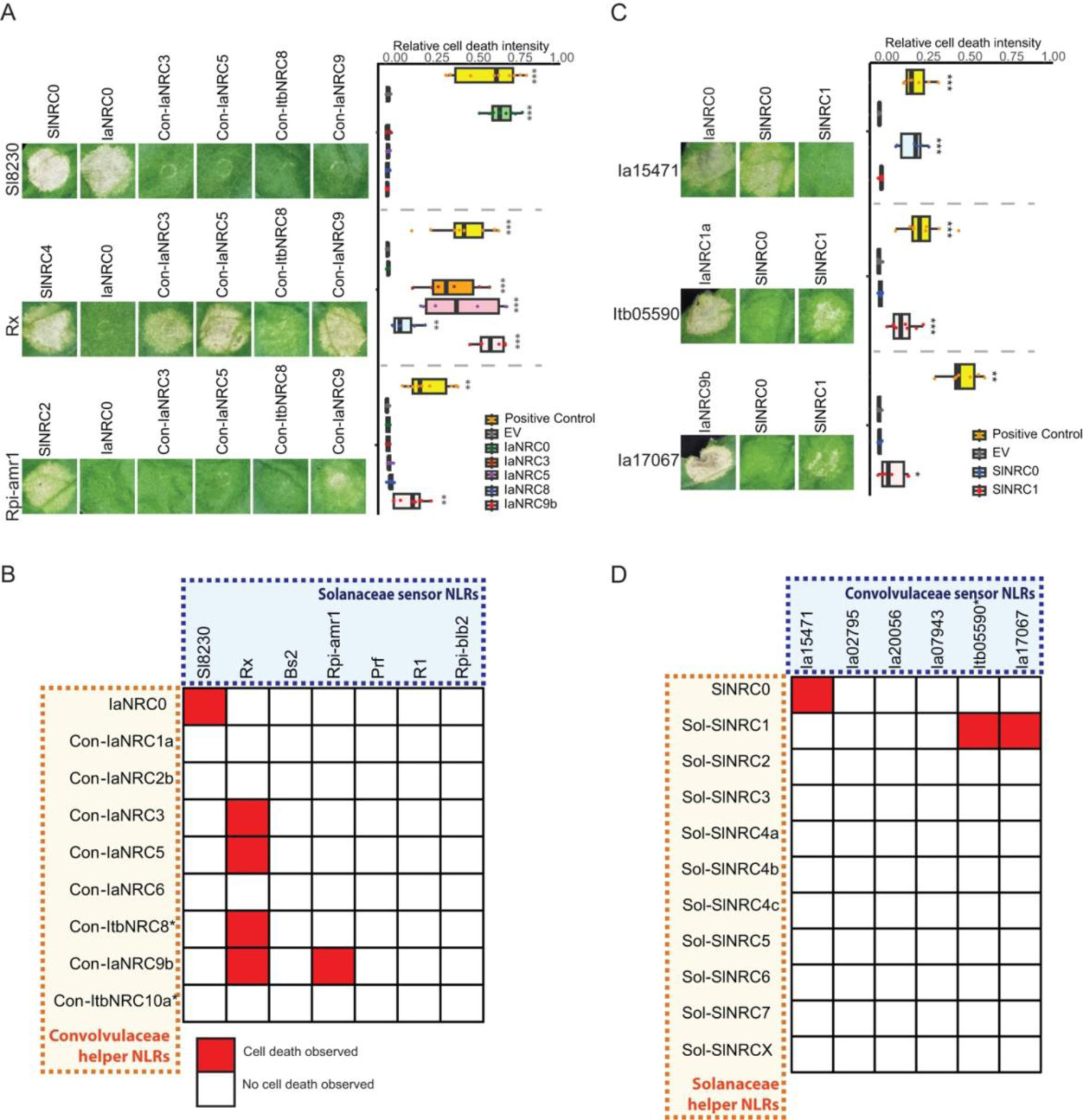
Sensor and helper NLRs in Solanaceae and Convolvulaceae exhibit a degree of cross-compatibility between these two plant families. **A)** Cell death assay results of solanaceous sensor NLRs (Sl8230, Rx, and Rpi-amr1) co-expressed with water spinach NRC helpers (IaNRC0, Con-IaNRC3, Con-IaNRC5, Con-IaNRC8, Con-IaNRC9) in *N. benthamiana* at 5 dpi. All sensor NLRs carried the MHD motif (D to V) mutation. The dot plot represents cell death quantification analysed by UVP ChemStudio PLUS. The bold line in the boxplots represents the medium, the box edges represent the 25th and 75th percentiles, and the whiskers extend to the most extreme data points no more than 1.5x of the interquartile range. Statistical differences were examined by paired Wilcoxon signed rank test. **B)** Matrix of cell death assays for solanaceous NRC-dependent sensor NLRs co-expressed with water spinach NRC helper NLRs, including information obtained from Supplemental Fig. S11 and S12. **C)** Cell death assay results of water spinach sensor NLRs (Ia15471, Ia04375, Ia14342, and Ia17067) co-expressed with solanaceous NRC helpers in *N. benthamiana* at 5 dpi. All sensor NLRs carried the MHD motif (D to V) mutation. The dot plot represents cell death quantification analysed by UVP ChemStudio PLUS. Statistical differences among the samples were analysed with Tukey’s HSD test (p < 0.05) for each sensor NLR independently. **D)** Matrix of cell death assays for solanaceous NRC-dependent sensor NLRs co-expressed with water spinach NRC helper NLRs, including information obtained from Supplemental Fig. S13 and S14.

## Discussion

The NRC network plays an essential role in disease resistance to multiple pathogens of solanaceous plants (Wu et al. 2017). Nevertheless, functional studies of the NRC network beyond solanaceous plants remained limited. We explored the evolutionary diversity of NRC networks across lineages of angiosperms, with a particular focus on the three major lineages of asterids (Ericales, campanulids, and lamiids). These lineages displayed distinct hierarchical structures within their respective NRC networks (Fig. 8). Ericales represents one of the early branches of asterids, predating the divergence of campanulids and lamiids. In our study, tea (*C. sinensis*) served as a representative of Ericales and possessed a relatively simple NRC network. This network involves the transmission of signals from NRC0-Eri-dependent sensor NLRs through NRC0-Eri to induce cell death. Since the three examined Ericales species contained few NRC superclade members, the NRC network in Ericales has likely undergone limited changes over millions of years of evolution (Fig. 8). In campanulids, we identified two partially redundant NRC nodes: NRC0 which is present in both Apiales and Asterales, and NRC0-Ast which exists exclusively in Asterales (Fig. 8). We speculate that NRC0-Ast may have either emerged early in campanulids and subsequently been lost in Apiales or arose via duplication of the ancestral NRC0 after the divergence of Asterales and Apiales. The NRC superclade expanded and diversified significantly in most lamiids (Fig. 1, Fig. 5, and Fig. 8). Apart from NRC0 which is very conserved, the phylogenetic tree shows distinct lineage-specific clustering patterns of the NRC family members. This specific diversification pattern appears to be unique for each plant family, implying that the lineage-specific NRC network is likely largely conserved at the family level (Fig. 5, Fig. 8, and Supplemental Fig S9). Our results are consistent with the view that the NRC superclade originated from an ancestral NRC sensor-helper pair, and then differentially expanded across plant lineages into complicated immune networks (Wu et al. 2017; Sakai et al. 2023).

**Figure 8.**
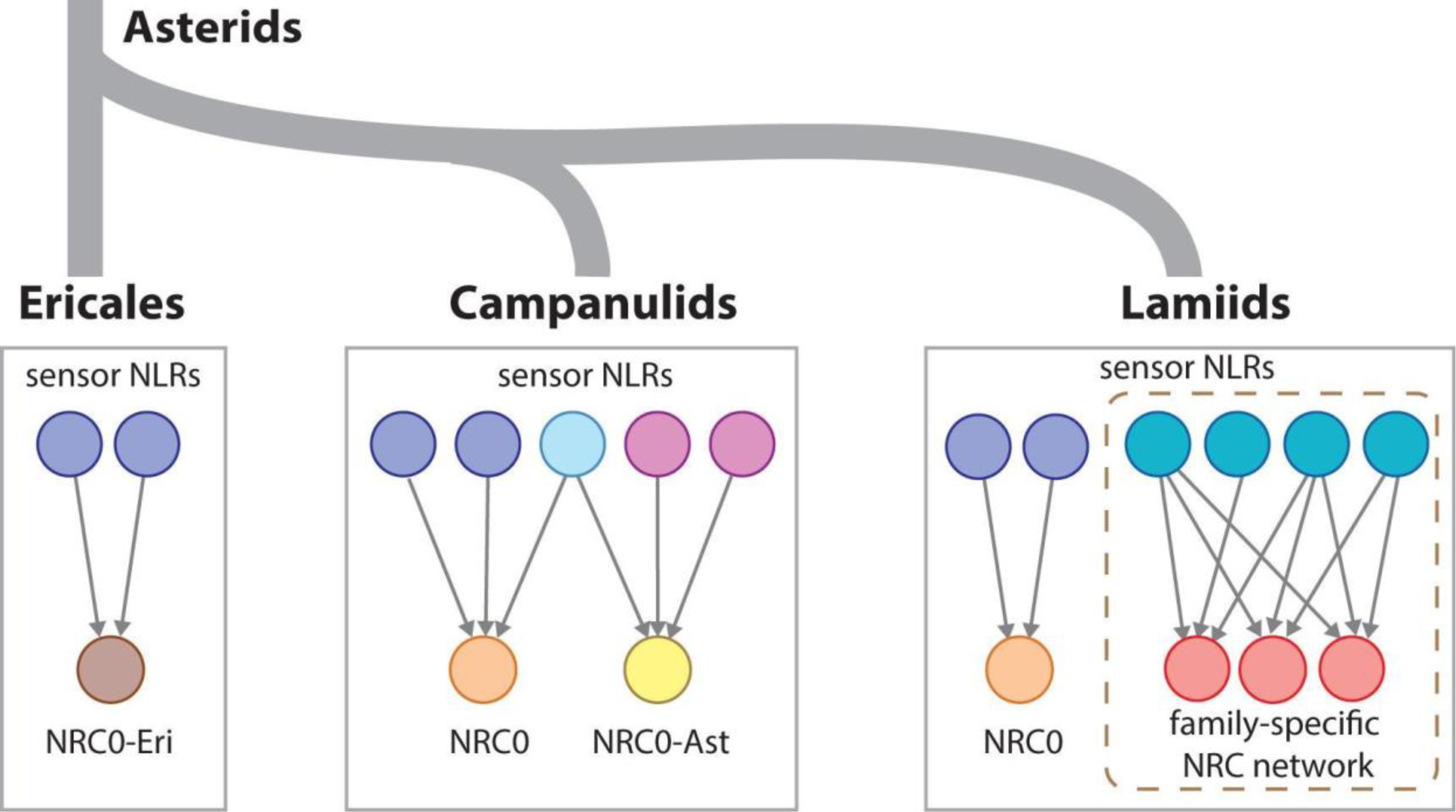
Evolution of the hierarchical structure of the NRC network in asterids. Ericales encode a simple NRC network where few sensor NLRs signal through NRC0-Eri to induce cell death. Campanulids show duplicated NRC nodes where some NRC-S signal through NRC0, some NRC-S signal through NRC0-Ast, and some signal through both NRCs to induce cell death. In addition to NRC0, lamiids display massively expanded and complex NRC networks that are likely plant family-specific.

We proposed that the ancestral state of the NRC network of superasterids (asterids together with Caryophyllales) is likely a sensor-helper cluster resembling the NRC0-H and NRC0-S described here and in Sakai et al., (2023) (Wu et al., 2017). Interestingly, the NRC-H identified in sugar beet (Caryophyllales) does not cluster with the NRC0 clades, which consists of NRC0/NRC0-Eri/NRC0-Ast, based on phylogenetic analyses (Supplemental Fig. S2A). Therefore, the ancestral NRC0 may have emerged in asterids from an NRC network shared with Caryophyllales, but this ancestral NRC network was subsequently lost in asterids. Alternatively, the ancestral NRC0 may have appeared early in the ancestral superasterids and was later lost in Caryophyllales. More phylogenetic and functional analyses using the NRC0 sequences identified from basal lineages of superasterids/asterids should help to clarify the early evolutionary history of the NRC network. Furthermore, the degree to which the phylogenetically defined Caryophyllales NRC-Hs function as helper NLRs and their cross-compatibility with sensor NLRs from asterids are still yet to be determined.

Tandem gene and whole-genome duplication (WGD), genetic drift, relaxed selection during domestication, and adaptation to ecological niches have all been reported to influence the expansion and contraction of *NLR* genes (Han and Tsuda 2022). While the total number of protein-coding genes shows a weak correlation with the expansion of NLRs, it is even more weakly correlated with the expansion of the NRC superclade (Fig. 1B and Supplemental Fig. S1). *Capsicum annuum*, for example, has a large number of NRC superclade members despite having relatively few total protein-coding genes (Fig. 1A and Supplemental Data Set 2) (Seo et al. 2016). Conversely, ginseng has high total protein-coding gene numbers but few NRC superclade members (Fig. 1A and Supplemental Data Set 2). Selection pressures placed on each plant lineage in asterids may, therefore, have led to distinct expansion or contraction patterns of the NRC superclade. Furthermore, the evolutionary events leading to the expansion of the NRC superclade show no correlation to the overall genome size (Supplemental Fig. S1). Since the massive expansion of the NRC superclade is a unique feature observed in lamiids, we speculate that selection pressure in the ancestral lamiids species may have led to the initial expansion of the NRC network. This ancient expansion provided advantages for the species to survive and was further selected independently in the subsequent lamiids progenies. While most lamiids species encode large numbers of NRC superclade members, *E. ulmoides* stands out as the only lamiids species analysed that does not have NLRs belonging to the NRC superclade (Fig. 1A). Although we have not yet been able to exclude the possibility that this is due to the insufficient quality of the assembled genome, additional genome information from related species may help address whether early gene loss events led to the absence of the NRC superclade in this species. Moreover, the identification of NLRs and overall protein content is influenced by the quality of both assembly and annotation. Future improvements in assembly and annotation should help to further verify the expansion and contraction of the NRC superclade in the species analysed here. Notably, 89% of NLRs predicted from the parasitic plant *S. asiatica* were classified within the NRC superclade, whereas *Cuscuta* spp., and many other parasitic plants, have undergone substantial loss of their NLRs (Fig. 1A) (Baggs et al. 2020; Liu et al. 2021). It would be interesting to understand whether and how the unique lifestyle of these and other parasitic plants may have differentially shaped the overall evolution of plant immunity, including the NRC superclade.

NLR genes are frequently located within gene clusters which can serve as a repository of genetic variation. Most of these clusters are results of tandem duplication leading to homogeneous NLR clusters, while heterogenous clusters are less prominent (Zhou et al. 2004; Meyers et al. 2005; Jupe et al. 2012). NRC0 stands out as the most conserved NRC homolog among asterids (Sakai et al., 2023). Furthermore, the NRC0 sensor-helper NLR gene cluster was identified in several species, likely representing the ancestral state of the asterids NRC network (Sakai et al., 2023). Based on the observation that most of the members in the NRC network do not remain as sensor-helper NLR gene clusters, we proposed that breaking the physical linkage was critical to enable the independent evolution of both sensor and helper NLRs. In Ericales, most of the sensor NLRs of the NRC superclade appeared to be truncated, and only those linked to NRC0-Eri retained full-length NLR sequence signatures (Fig. 2 and Supplemental Fig. S2). A similar phenomenon was observed in carrots (campanulids), where only the sensor NLRs that were closely linked to NRC0 remained functional (Supplemental Fig. S3). In lettuce, the three sensor NLRs that signal through LsNRC0-Ast are located on the same chromosome as LsNRC0-Ast, whereas LsNRC0 is physically disconnected from the sensor NLRs (Supplemental Fig. S4). The group of sensor NLRs that signal through LsNRC0 show longer branch lengths in the phylogenetic tree compared to the NRC0-Ast-dependent sensor NLRs group (Supplemental Fig. S4A). This supports the hypothesis that, after becoming physically unlinked from their helper NLR partners, the sensor NLRs are more prone to accumulating mutations, leading to diversification or non-functionalisation (Baumgarten et al. 2003; Leister 2004). In water spinach, we identified only a few sensor-helper NLR clusters in addition to the NRC0 cluster (Supplemental Fig. S6). Similar to what was observed in solanaceous crops, most of the water spinach NRC superclade members are dispersed on different chromosomes (Supplemental Fig. S6) (Seo et al. 2016; Wu et al. 2017). These results suggest that the expansion of the NRC superclade coincides with the transposition and tandem duplication of both sensor and helper NLR genes independently. Transposable elements, particularly the long terminal repeat retrotransposons, were shown to contribute to the tandem duplication and transposition of NLR genes in plant genomes (Wei et al. 2002; Kim et al. 2017; Seidl and Thomma 2017; Hao et al. 2023). Whether the associations with transposable elements correlate with the expansion of the NRC superclade in different plant lineages requires further investigation.

The asterids NRC superclade likely originated from an ancient sensor-helper NLR cluster that contained a one-to-one or two-to-one ratio of sensor and helper NLRs (Wu et al. 2017; Sakai et al. 2023). Our results indicate that NRC-S duplicated more times than NRC-Hs in most species analysed and generally showed longer branch lengths than that of helper NLRs in the phylogenetic trees (Fig. 1C, Fig. 2A, Fig. 3A, and Fig. 4A). This is consistent with the view that sensor NLRs are often under high dynamic and balancing selection, whereas helper NLRs show slower evolution rates and remain functionally conserved (Seo et al. 2016; Stam et al. 2019; Shimizu et al. 2022). Furthermore, an expanded and diversified sensor NLR repertoire offers a higher potential for conferring resistance to pathogens (Barragan and Weigel 2021). In campanulids, where the NRC superclade is not as extensively expanded, certain species such as *S. rebaudiana* encode a high sensor-to-helper ratio (Fig. 1C). This is perhaps due to several tandem duplication and transposition events that occurred to NRC-S but not to NRC-H in these species. Although sensor NLRs are generally highly expanded in lamiids, leading to a high sensor-to-helper ratio, *I. cairica*, *I. aquatica*, and *N. benthamiana* are among the few species in which the sensor-to-helper ratio remains close to the ancestral ratio (Fig. 1C). As NLR contraction events have been reported in lineages of Brassicaceae and Apiaceae species (Shao et al. 2016; Zhang et al. 2016; Liang and Dong 2023), one possible explanation is that NLR contraction contributes to the reduction of the sensor-to-helper ratio, with many of sensor NLRs being lost during evolution. Additional macroevolutionary analyses with diverse plant genomes may provide insights into how NLR expansion and contraction events influence sensor-helper NLR ratios and provide information on the factors that drive NLR contraction.

NRC0 stands out as the most conserved NRC across different plant lineages (Sakai et al. 2023). We found two other NRC0 subclades, namely NRC0-Eri and NRC0-Ast, which show partial interchangeability with the NRC0 clade defined by Sakai et al. (2023) (Fig. 6). The three NRC0 subclades show relatively short branch lengths, indicating that NRC0 homologs have not undergone significant diversification since its split from the ancestral species (Fig. 5). Recent macroevolutionary studies of ZAR1 homologs revealed that ZAR1 is the most conserved NLR across angiosperms and evolved to function with partnered RLCKs early in its evolutionary history (Adachi et al. 2023b). While RLCKs rapidly diversified to keep pace with fast-evolving effectors, ZAR1 experienced relatively limited expansion and duplication (Adachi et al. 2023b). Whether the conservation of NRC0 subclades is due to its critical role in immunity or simply a lack of pathogen pressure remains to be investigated. The phylogenetic tree of the NRC family in lamiids indicated that, apart from the highly conserved NRC0s, there are no clear orthologous NRCs across different plant families (Fig. 4B, and Fig. 5A). Nevertheless, some NRC-S from solanaceous plants can activate some *Ipomoea* NRCs to induce cell death, and some NRC-S from *Ipomoea* are capable of inducing cell death through SlNRC1 (Fig. 7). Therefore, while NRCs function as helper NLRs that mediate cell death for NRC-S, this group of genes has diversified to an extent that functional orthologs (such as NRC0) are rarely found in plants from different families. Despite this, compatibility between sensor and helper NLR across different plant families has been observed (e.g. between Solanaceae and Convolvulaceae). Further studies focusing on understanding the molecular mechanisms underpinning these interactions may help elucidate the critical changes that determine compatibility between NRCs and their matching sensor NLRs.

We tested many NRC-S and NRC-H combinations using the heterologous expression system in *N. benthamiana*. Although many of the NRC-S/H combinations induced cell death, some combinations did not trigger cell death in *N. benthamiana*. In particular, NRC0-Eri was the only NRC-H group identified in tea and carrot, but not all the NRC-S of tea and carrot can signal through the tested NRC0-Eri. Similar phenomena were observed with other NRC-S cloned from *L. sativa* and *I. aquatica* that did not trigger cell death with any NRC-H tested. While some of these NRC-S may have accumulated loss-of-function mutations, we could not rule out that additional putative NRC-H members from these species were missing in the annotated genome, or an alternative mutation (such as H to R rather than D to V mutation in the MHD motif) in the NLR is required to trigger cell death, such as the case for NbNRC2 (Derevnina et al., 2021). It is also possible that an essential component for these NRC-S/H combinations to function is lacking in *N. benthamiana*. Further studies with improved genome information, alternative mutations, or expression in the original plant species may help to address this issue.

Our findings reveal the overall diversity and hierarchical structure of NRC networks in plants belonging to different lineages of asterids. Except for a small number of NRC-H that can cooperate with NRC-S from different plant lineages, the majority of NRC-H do not function with NRC-S from divergent plant species. Knowledge of sensor-helper compatibility may be particularly useful for overcoming restricted taxonomic functionality, a challenge when transferring resistance genes across distantly related plants (Tai et al. 1999; Narusaka et al. 2013) In the cases of NLRs belonging to the NRC network, it may be necessary to transfer both the matching sensor and helper NLRs together. The recent discovery that CaRpi-blb2 specifically signals through the pepper helper NLRs CaNRC8 and CaNRC9 further emphasizes the importance of understanding the commonalities and differences in the immunity networks of various related species (Oh et al. 2023). Among the *Ipomoea* species, *I. trifida, I. tiloba,* and *I. batatas* encode high numbers of overall NLRs in their genomes. Given that the majority of NLRs in these three species belong to the NRC superclade, delving into the NRC network of *Ipomoea* species might offer opportunities to utilize their NLRome as a resource for conferring disease resistance against a variety of pathogens. Further investigations into the evolution of the NRC network, along with validation using NRC-S that confer resistance in combination with their corresponding NRC-H, could increase the likelihood of successfully transferring disease resistance across distantly related crops.

## Materials and methods

### Prediction and phylogenetic analyses of NLRs

The annotated protein sequences of selected plant species used in this study were downloaded from public databases (Supplemental Data Set 1). NLR prediction and phylogenetic classification were performed according to previously established pipelines (Jupe et al. 2012; Andolfo et al. 2014). To identify the proteins containing NLR features, sequences were scanned with MAST in MEME v 5.4.0 using 20 previously defined motifs with default parameters (Bailey et al. 2009; Jupe et al. 2012). The extracted sequences were then aligned using MAFFT version 7 with the G-INS-1 progressive method and default settings (Katoh et al. 2019). The aligned amino acid sequences were imported into MEGA 7 for manual trimming leaving only the NB-ARC domain. The NB-ARC domains with at least three of the four major motifs (P-loop, GLPL, Kinase2 and MHD) were considered intact. Sequences with truncated NB-ARC domains were excluded from further analyses. The remaining alignments are usually around 250 amino acids in length depending on the sequences included. Phylogenetic trees were constructed in MEGA 7 using Maximum-likelihood phylogenetic analyses with the evolutionary model JTT+G+I and 500 bootstrap tests (Kumar et al. 2016). The number of bootstrap tests required were conducted using a convergence test in RAxML (Stamatakis 2014). Phylogenetic trees were further processed and visualized using FigTree and iTOL (Rambaut 2018; Letunic and Bork 2021). NLR sequences from tomato were included as references for the identification of the NRC superclade. The species tree of angiosperms in Fig. 1 was modified from the published phylogenetic tree (The Angiosperm Phylogeny Group 2016). All datasets used for phylogenetic analyses are in Supplemental Files 1.

### Detection of positive selection

Full-length nucleotide sequences of the NRC family were analysed using MEGA 7. The nucleotide sequences were translated into protein sequences and subsequently aligned using ClustalW. Gaps in the alignment were manually removed. The resulting trimmed amino acid sequences of the NRC family were subjected to phylogenetic analyses. Phylogenetic trees were constructed using Maximum-likelihood phylogenetic analyses with the evolutionary model JTT+G+I and 200 bootstrap tests. We then generated codon-based alignment of the NRC family using the amino acid sequences alignment mentioned above. Along with the phylogenetic tree, codon-based alignment of the NRC family was subjected to aBSREL (An adaptive branch-site REL test for episodic diversification) using the HyPhy (Hypothesis testing using Phylogenies) package on the Datamonkey Adaptive Evolution Server (https://www.datamonkey.org/) (Smith et al. 2015; Weaver et al. 2018; Kosakovsky Pond et al. 2020) SlNRC0-dependent sensor NLR Sl8230 (Solyc10g008230) was included as an outgroup.

### RNA and DNA isolation

Materials of carrot (*Daucus carota*), kiwifruits (*Actinidia chinensis*), tea (*Camelia sinensis*), lettuce (*Lactuca sativa*), and water spinach (*Ipomoea aquatica*) were obtained from local nurseries or grocery stores. Littlebell (*Ipomoea triloba*) was collected from the campus of Academia Sinica (Nankang, Taipei, Taiwan) and confirmed by Sanger sequencing using ITK and MatK primers. DNA was extracted from young seedlings or mature leaves using DNeasy Plant Mini Kit (Qiagen). RNA was extracted using Plant Total RNA Mini Kits (VIOGENE). Synthesis of cDNA was performed using SuperScript™ III Reverse Transcriptase (Invitrogen) following the manufacturer’s instructions.

### Cloning of NLR genes

Full-length NLRs were amplified from genomic DNA or cDNA with primers designed based on the available genome sequences. The PCR amplicons were cloned into the pAGM9121 (Addgene plasmid #51833), pICH41308 (Addgene plasmid #47998) using Golden Gate Cloning (Weber et al. 2011) or pTA in T&A™ Cloning Kit from Yeastern Biotech (Taiwan). The putative sensor NLRs of tea (Cs0021741 and Cs0024074), and *I. aquatica* NRC6 were synthesized in pUC57-Kan as MoClo level 0 modules using the service provided by SynBio Technologies (New Jersey, USA).

Mutations (D to V) of the MHD motif were carried out using inverse PCR with primers containing AarI (NEB) or BsmBI (NEB) enzyme sites for digestion followed by ligation using T4 DNA ligase (Invitrogen). The constructs were then subcloned into binary vector pICSL86922OD using BsaI (NEB) and T4 DNA ligase (Invitrogen). pICSL86922OD was kindly provided by Mark Youles (The Sainsbury Laboratory, UK) (Addgene plasmid # 86181). The NLR gene expression constructs were then transformed into *Agrobacterium tumefaciens* (GV3101) using electroporation. The NLR gene sequences and cloning primers used in this research are listed in Supplemental Data Set 3 and Supplemental Data Set 5. Several previously described NRC-dependent solanaceous NLRs, including R1, Rx, Rpi-amr1, Rpi-blb2, Bs2, and Prf (Pto), and the corresponding effectors, including AVR1, CP, AVRamr1, AVRblb2, AvrBs2, and AvrPto were used in the cell death assays together with Con-IaNRCs (Wu et al. 2017; Witek et al. 2021).

### Agroinfiltration and quantification of cell death in *N. benthamiana* leaves

Transient expression of NLRs were performed on 4-week-old *N. benthamiana* leaves (WT or *nrc2/3/4*_KO) with *A. tumefaciens* (GV3101) carrying the indicated expression constructs (Wu et al. 2017; Witek et al. 2021). *A. tumefaciens* suspensions in infiltration buffer (10 mM MES, 10 mM MgCl2, and 150 µM acetosyringone, pH 5.6) were adjusted to suitable OD_600_ and then infiltrated into *N. benthamiana* leaves using needleless syringes. The agro-infiltrated plants were kept in a walk-in growth chamber (temperature 24-26°C, humidity 45–65% and 16/8 hr light/dark cycle) for 5 days before imaging and autofluorescence-based cell death quantification using UVP ChemStudio PLUS (Analytik Jena). Raw cell death autofluorescence images were acquired using blue LED light for excitation and the FITC filter (513 - 557 nm) as the emission filter. Areas showing stronger visual cell death generally produce stronger autofluorescence at this wavelength range. The exposure time was adjusted to 10 seconds to avoid saturation of the autofluorescence signal. The mean signal intensity was calculated using VisionWorks v.11.2 software by manually selecting the infiltrated areas and subtracting the background signal intensity. The mean signal intensity value was further normalized with the maximum intensity (65535) to obtain the relative intensity of cell death. Results of cell death assays were presented as relative cell death intensity from at least 6 technical replicates. In some cases, co-expression of NRC-H and NRC-S induces weak and variable cell death. Quantification using autofluorescence-based analyses may not yield statistically significant results due to the variation observed for the weak phenotypes. We classified these as weak cell death to distinguish them from those that completely lacked cell death (e.g. where no visible cell death was observed).

### Statistical analysis of cell death assays

Statistical analysis was performed using R (v4.3.3). Nonparametric Wilcoxon signed rank tests were conducted to compare the negative controls (EV) and the tested groups using the rstatix package in RStudio (Kassambara 2023). Statistical significance was defined as p < 0.05 and represented in figures as follows: *, p < 0.05; **, p < 0.01; ***, p < 0.001. Detailed statistical data are provided in Supplementary Data Set 6.

## Supporting information

Supplemental data set S1-6

## Acknowledgements

We thank Dr. Sophien Kamoun (The Sainsbury Laboratory, UK), Dr. Hiroaki Adachi (Laboratory of Crop Evolution, Graduate School of Agriculture, Kyoto University) and Toshiyuki Sakai (Laboratory of Crop Evolution, Graduate School of Agriculture, Kyoto University) for valuable suggestions on the research and comments on the manuscript. We thank Dr. Tsai-Ming Lu (Institute of Cellular and Organismic Biology, Academia Sinica) for helping with phylogenetic analyses. We thank Mark Youles (SynBio, The Sainsbury Laboratory, UK) for sharing plasmids for molecular cloning. We also thank Dr. Chih-Horng Kuo, Dr. Lay-Sun Ma, and Dr. Ting-Ying Wu (Institute of Plant and Microbial Biology, Academia Sinica) for their suggestions on the experimental design and data presentation. This research is funded by the Institute of Plant and Microbial Biology, Academia Sinica. C.H.W. is funded by the 2030 Cross-Generation Young Scholars Program of the National Science and Technology Council, Taiwan (NSTC 112-2628-B-001-007). L.D. is funded by a National Institute of Agricultural Botany (NIAB) Fellowship. Research in the LD lab is supported by the British Society of Plant Pathology, the Gatsby Charitable Foundation and the Royal Society.

## Author Contributions

F.J.G. and C.H.W. designed the research. F.J.G. and C.Y.H. conducted the experiments. F.J.G. analysed the data. F.J.G., L.D., and C.H.W. wrote the manuscript.

## Notes

### Competing Interest Statement

The authors have declared no competing interest.

### Summary of Updates

Discussion updated; Figures 2,3,4,6,7 revised; Supplemental files updated.

