## Supplemental Figures 1-15 for "NRC immune receptor networks show diversified hierarchical genetic architecture across plant lineages"

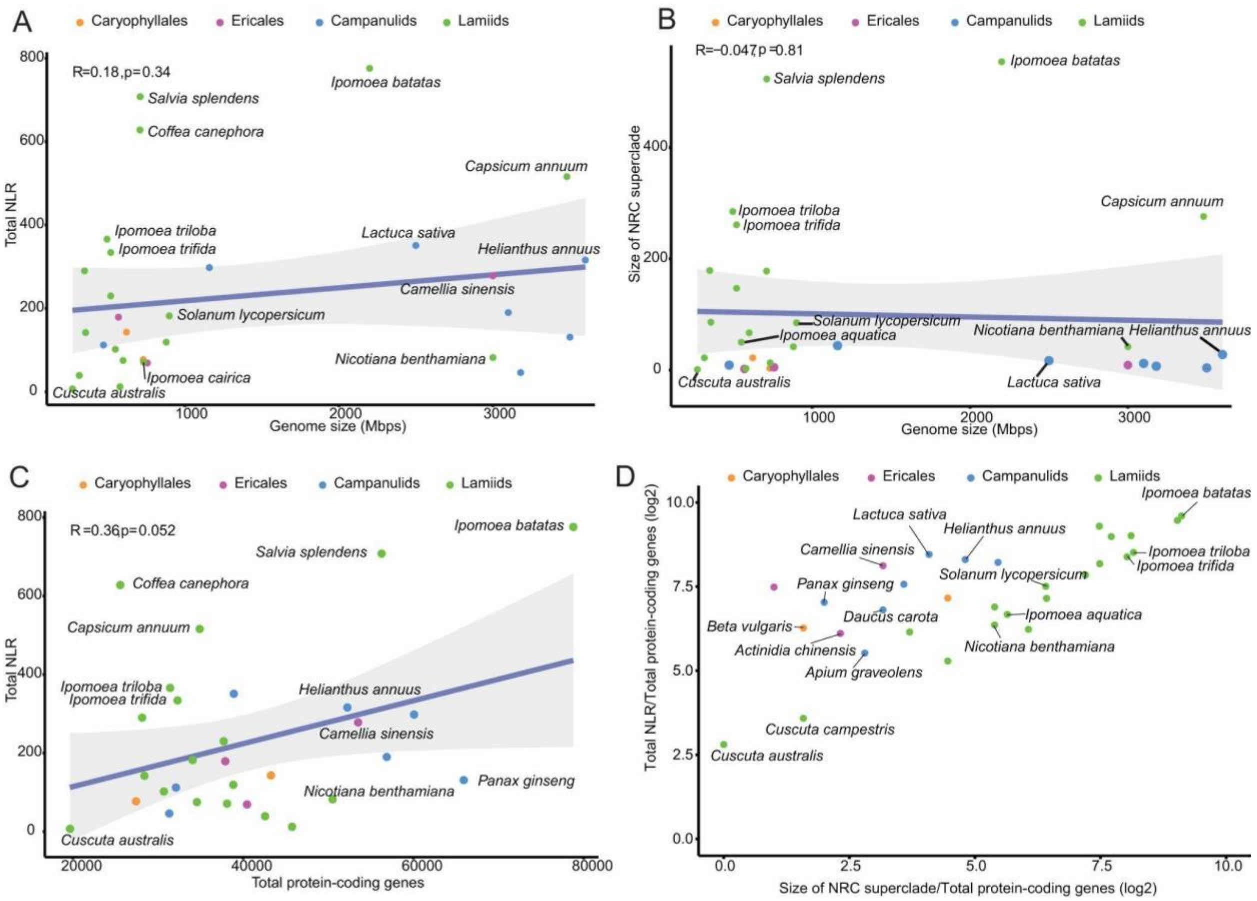


**Supplemental Fig. S1.** Correlation of the size of total NLRs or NRC superclade with total genome size or total protein-coding genes. **A)** Total NLR and genome size show weak correlations. **B)** The size of the NRC superclade and total protein-coding genes shows no correlation. **C)** Total NLR and total protein-coding gene showed a weak correlation. **D)** Lamiids plant species showed a higher size of NRC superclade from total NLR out of total protein-coding genes (log2). Plant species from different lineages indicated are labelled with different colours.

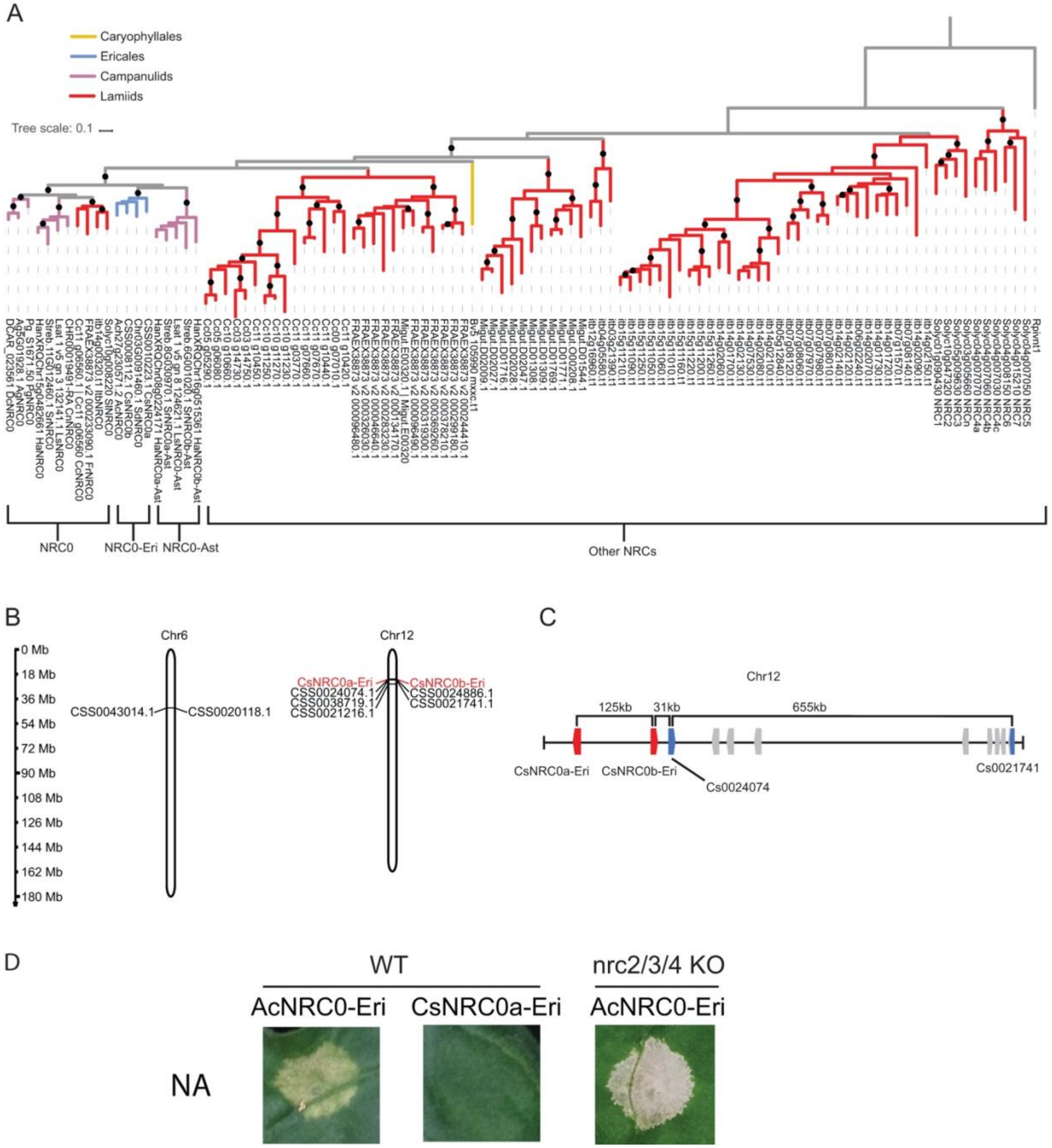


**Supplemental Fig. S2.** Tea NRC0-Eri and putative sensor NLRs are located in a gene cluster. **A)** Phylogenetic analysis of NRC helpers from Caryophyllales (*B. vulgaris*), Ericales, campanulids and lamiids. Nodes with bootstrap values over 70 are labelled with black dots. The scale bar indicates the evolutionary distance in amino acid substitution per site. **B)** The NRC superclade members of tea (*C. sinensis*) form gene clusters on chromosome 6 and chromosome 12. **C)** The NRC0 sensor and helper genes cluster together on chromosome 12. **D)** The kiwifruit NRC0 exhibited autoactivity when expressed in *N. benthamiana* leaves. "WT" represents Wild type *N. benthamiana* plants, and "nrc2/3/4 KO" represents *nrc2/nrc3/nrc4* triple knockout *N. benthamiana* plants.

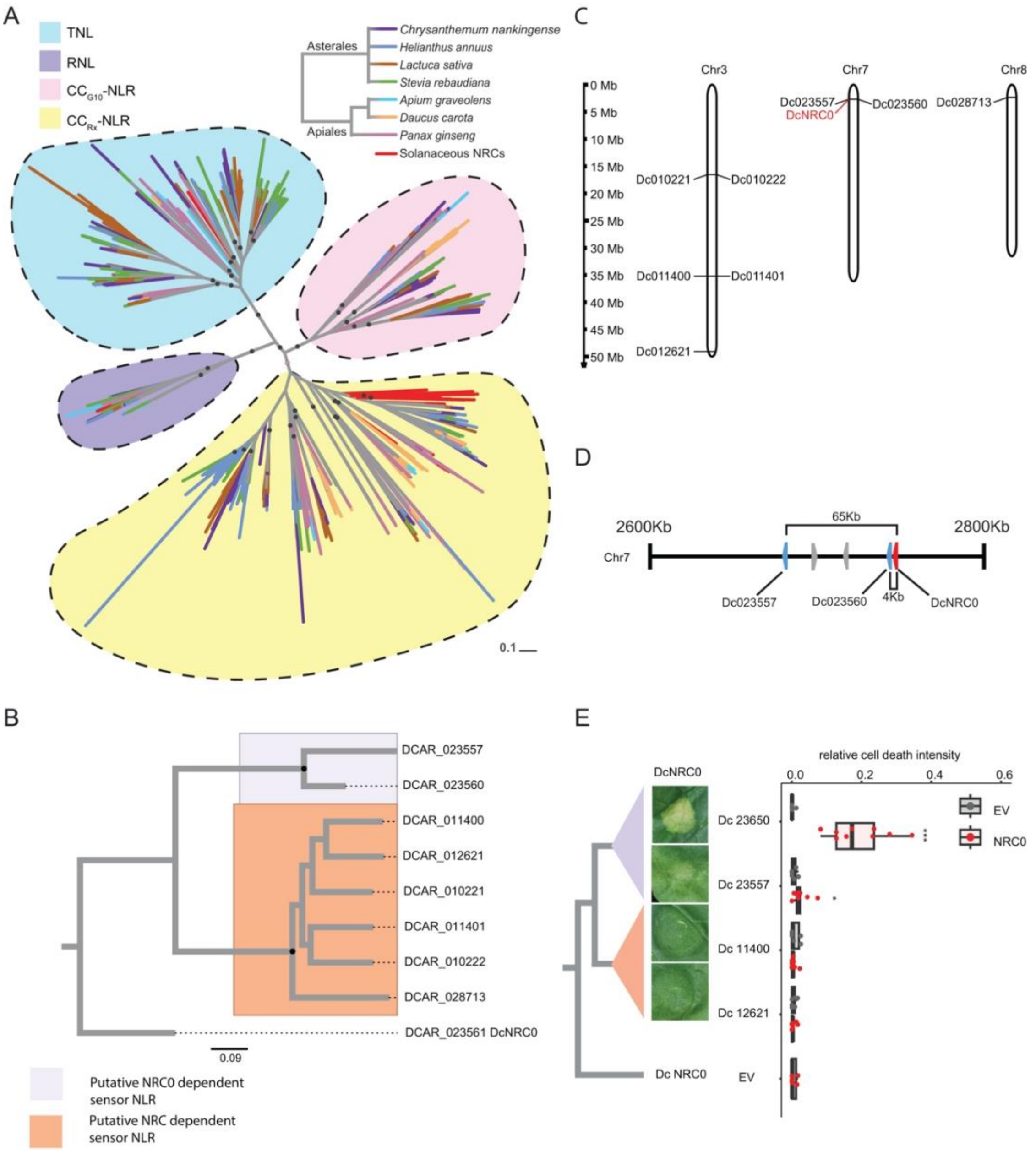


**Supplemental Fig. S3.** Carrot NRC0-dependent sensor NLRs induce cell death through the linked NRC0. **A)** Phylogenetic analysis of NLRs from 7 species of campanulids. Major nodes with bootstrap values over 70 are indicated with black dots. The scale bar indicates the evolutionary distance in amino acid substitution per site. **B)** The NRC superclade members of carrot (*D. carota*) on chromosome 3, chromosome 7, and chromosome 8. **C)** The NRC0 sensor and helper genes cluster on chromosome 7. **D)** Phylogenetic analysis of the NRC superclade of carrot. Major nodes with bootstrap values over 70 are indicated with black dots. **E)** Cell death assay results of NRC-dependent sensor NLRs co-expressed with the NRCs from carrot in *N. benthamiana* observed at 5 dpi. All sensor NLRs carried the MHD motif (D to V) mutation. The dot plot represents cell death quantification analysed by UVP ChemStudio PLUS. The bold line in the boxplots represents the medium, the box edges represent the 25th and 75th percentiles, and the whiskers extend to the most extreme data points no more than 1.5x of the interquartile range. Statistical differences were examined by paired Wilcoxon signed rank test (* = $p < 0.05$, and *** = $p < 0.001$).

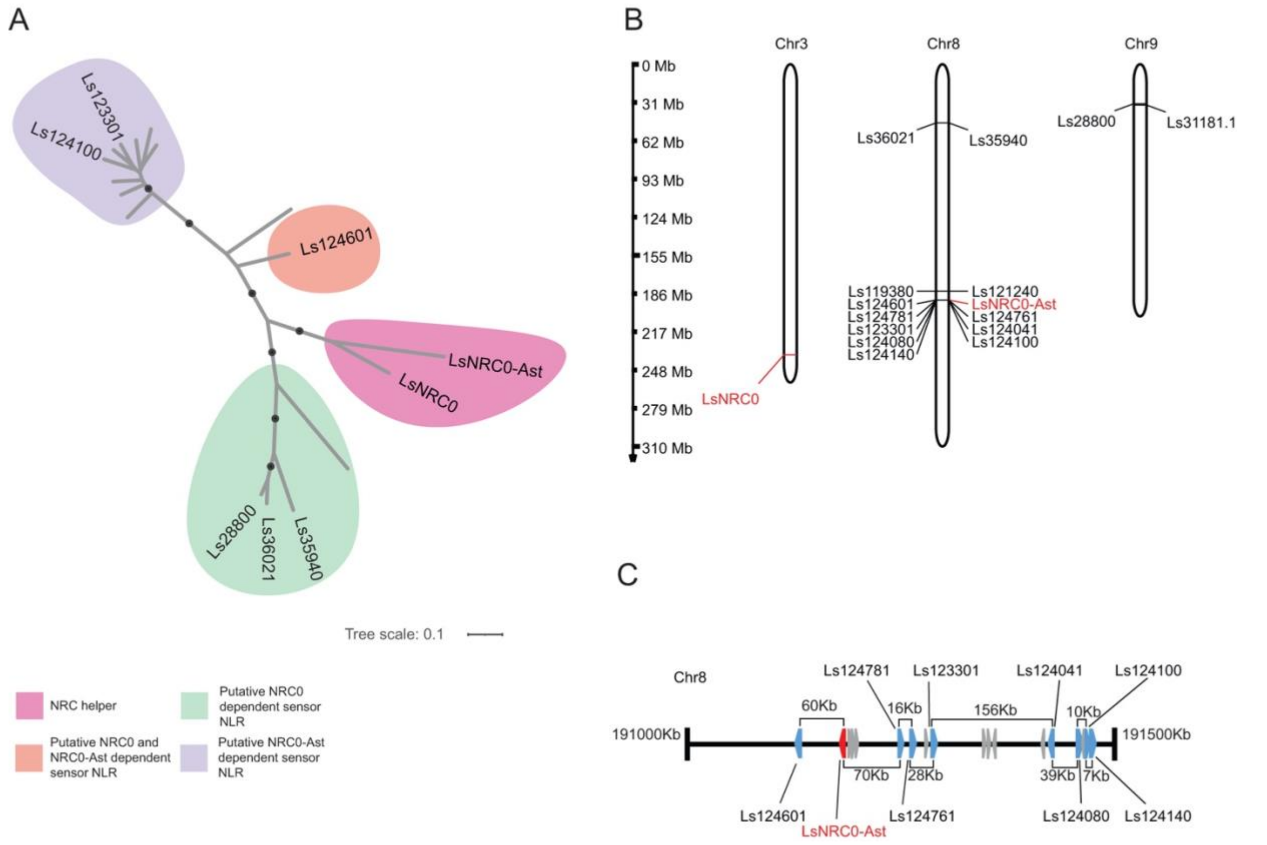


**Supplemental Fig. S4.** Phylogenetic analysis and gene cluster of the NRC superclade in lettuce. **A)** Phylogenetic analysis of the NRC superclade of *L. sativa*. Major nodes with bootstrap values over 70 are indicated with black dots. Different colors indicate the NRC helper NLR clade and NRC sensor NLR clades with different putative NRC dependency based on the results obtained in Fig. 3C. The scale bar indicates the evolutionary distance in amino acid substitution per site. **B)** The NRC superclade members of lettuce (*L. sativa*) on chromosome 3, chromosome 8, and chromosome 9. **C)** The NRC0 sensor (blue) and helper (red) genes cluster on chromosome 8.

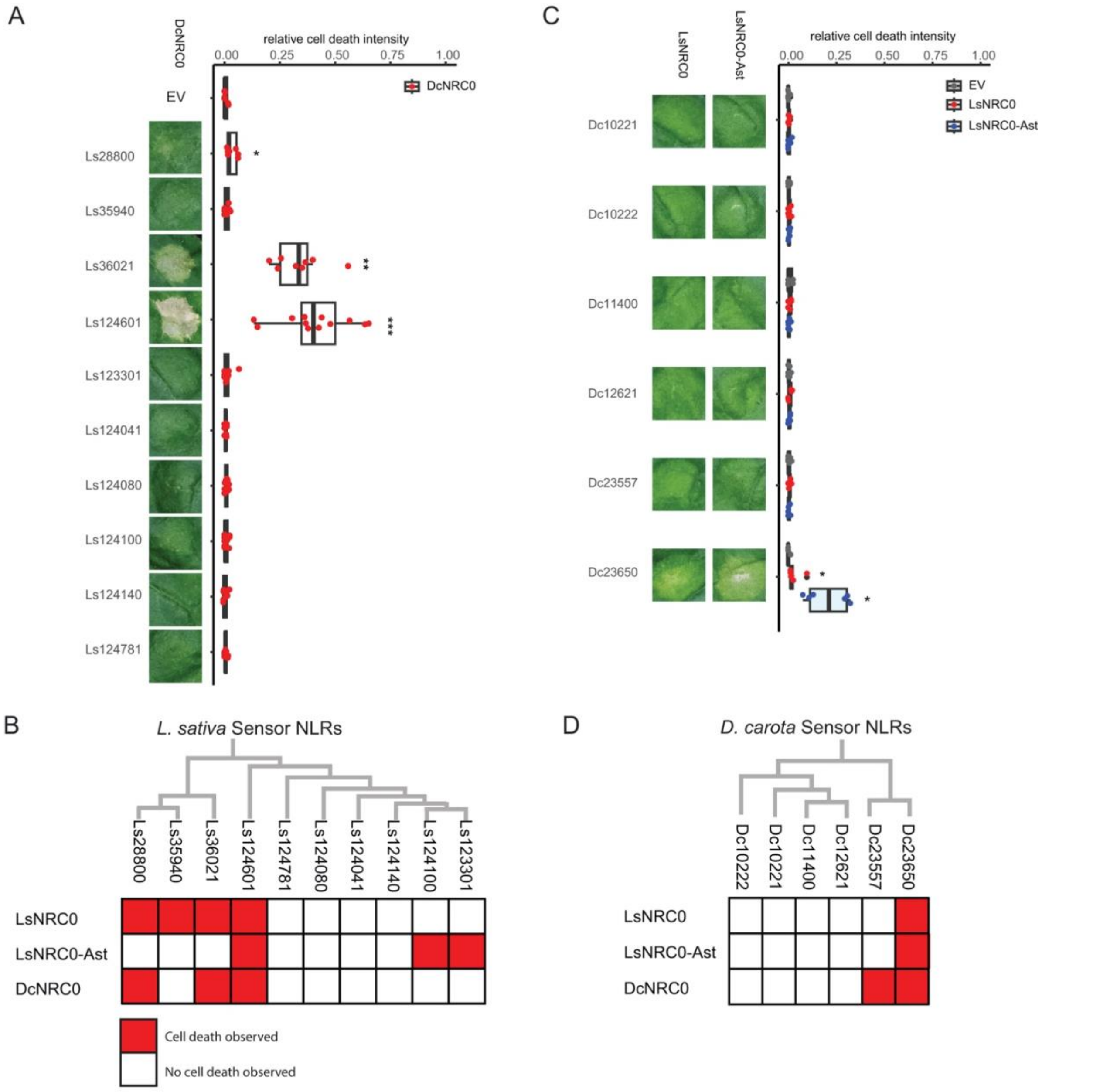


**Supplemental Fig. S5.** Lettuce LsNRC0 and LsNRC0-Ast are partially interchangeable with carrot DcNRC0. **A)** Cell death assay results of lettuce putative NRC-dependent sensor NLRs co-expressed with carrot NRC0 in *N. benthamiana* observed at 5 dpi. **B)** Matrix of cell death assays for lettuce sensor NLRs co-expressed with lettuce and carrot NRCs, including information obtained in Fig. 3. **C)** Cell death assay results of carrot putative NRC-dependent sensor NLRs co-expressed with lettuce NRCs in *N. benthamiana* observed at 5 dpi. All sensor NLRs carried the MHD motif (D to V) mutation. For (A) and (C), the dot plot represents cell death quantification analysed by UVP ChemStudio PLUS. The bold line in the boxplots represents the medium, the box edges represent the 25th and 75th percentiles, and the whiskers extend to the most extreme data points no more than 1.5x of the interquartile range. Statistical differences were examined by paired Wilcoxon signed rank test (* = $p < 0.05$, ** = $p < 0.01$; *** = $p < 0.001$). **D)** Matrix of cell death assays for carrot sensor NLRs co-expressed with lettuce and carrot NRC families, including information obtained in Supplemental Fig. S3.

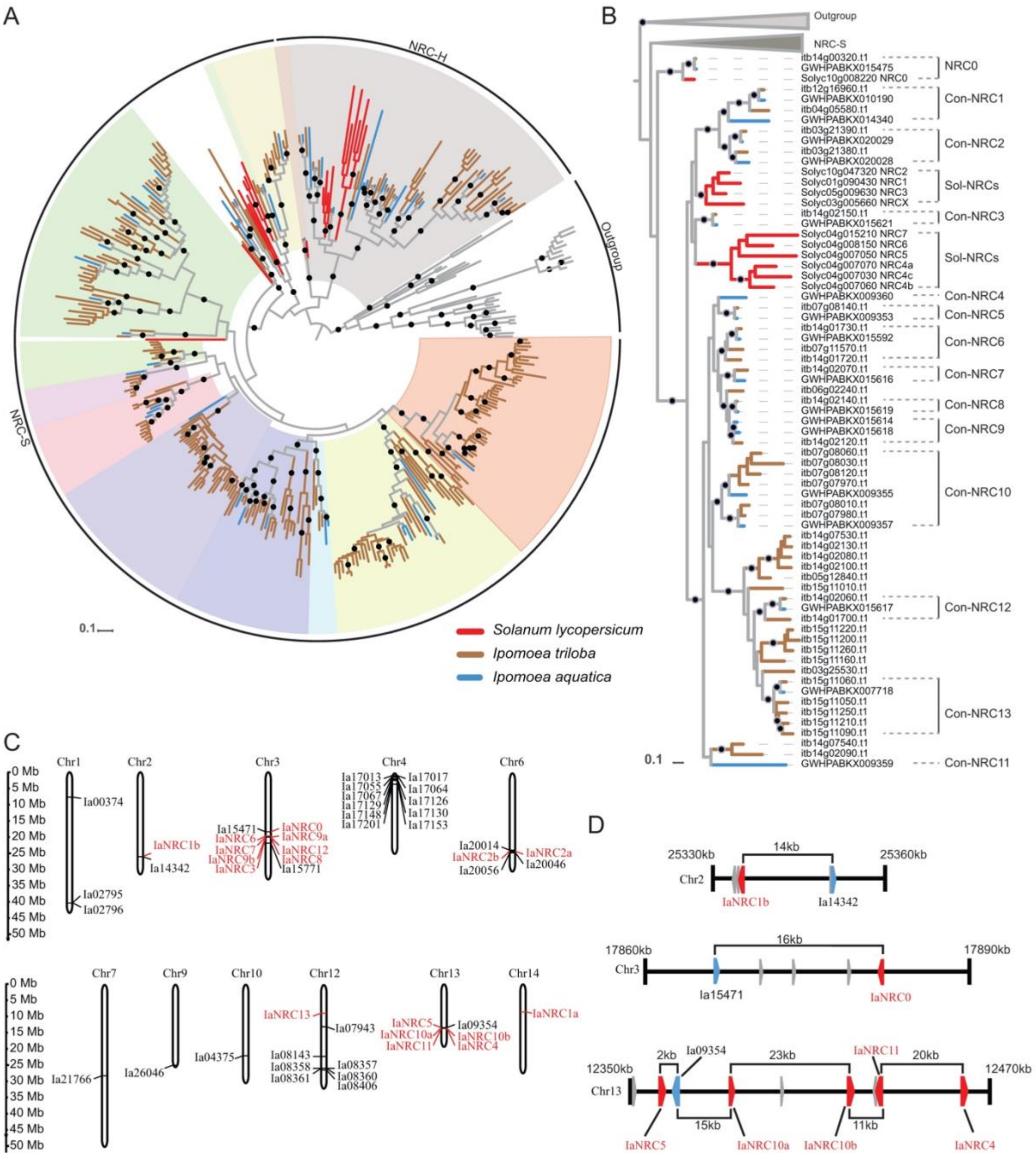


**Supplemental Fig. S6.** *I. aquatica* has a smaller and simpler NRC network compared to *I. triloba*. **A)** Phylogenetic analysis of the NRC superclade of tomato, *I. triloba*, and *I. aquatica*. Nodes with bootstrap values over 70 are labelled with black dots. Sequences belonging to different species are indicated with lines of different colours. Different well-supported NRC-H or NRC-S clades are labelled with different background colours. The scale bar indicates the evolutionary distance in amino acid substitution per site. **B**) Phylogenetic tree of NRC family from *I. aquatica* and *I. triloba*. The different Con-NRCs are defined based on the bootstrap supports in each phylogenetic group. The scale bar indicates the evolutionary distance in amino acid substitution per site. **C)** The distribution of NRC superclade members of water spinach (*I. aquatica*) on different chromosomes. **D)** The NRC sensor (blue) and helper (red) genes cluster on chromosome 2, chromosome 3, and chromosome 13.

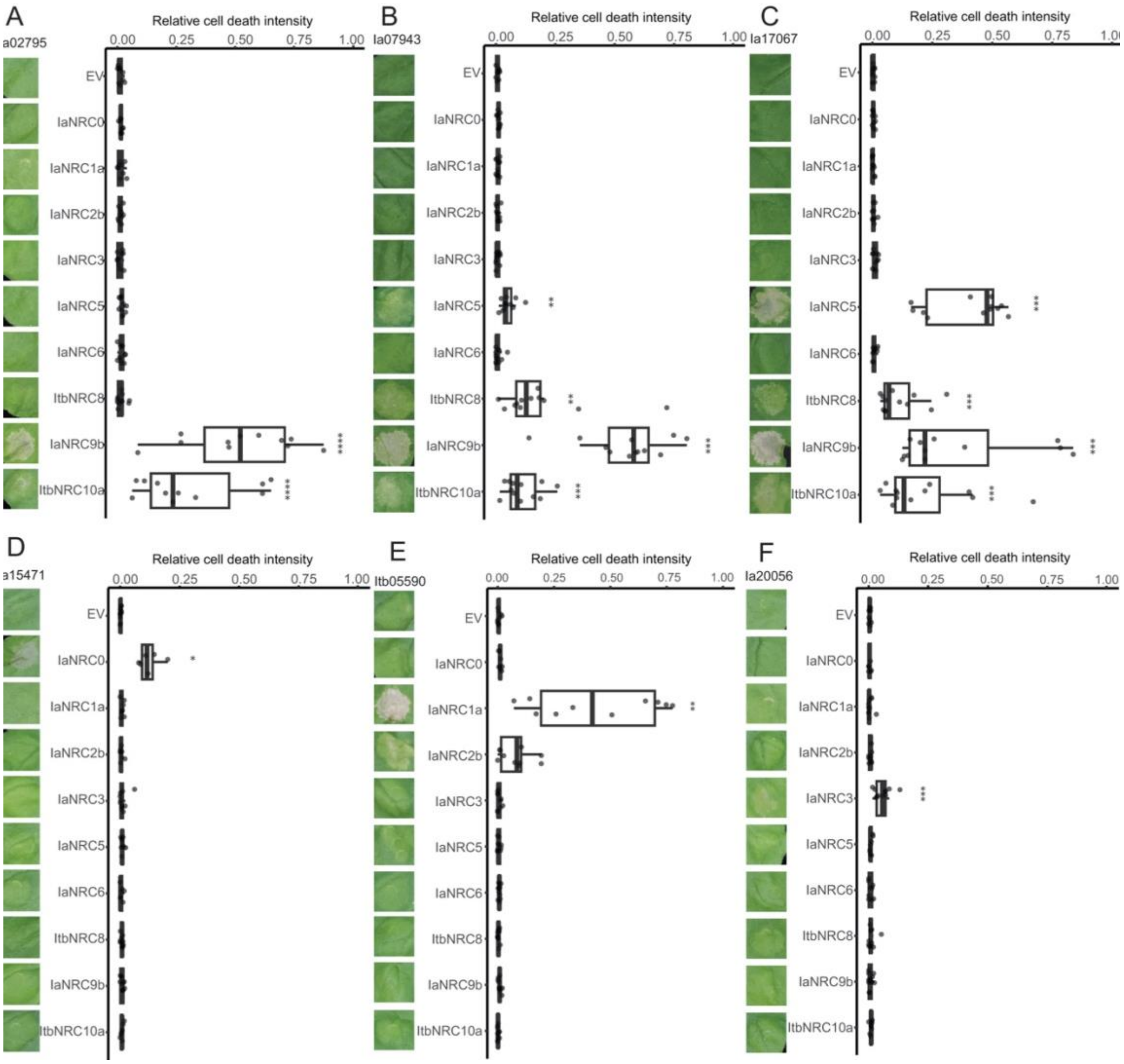


**Supplemental Fig. S7.** Cell death assay results of *I. aquatica* sensor NLRs co-expressed with Con-IaNRCs in *N. benthamiana* observed at 5 dpi. *I. aquatica* sensor NLRs **A)** Ia02795, **B)** Ia07943, **C)** Ia17067, **D)** Ia04375, **E)** Ia15471, **F)** Ia14342, and **G)** Ia20056 were made into autoactive (D to V mutation in the MHD motif) and then co-expressed with Con-IaNRCs in *N. benthamiana.* The dot plot represents the cell death quantification analysed by UVP. The bold line in the boxplots represents the medium, the box edges represent the 25th and 75th percentiles, and the whiskers extend to the most extreme data points no more than 1.5x of the interquartile range. Statistical differences were examined by paired Wilcoxon signed rank test (* = p < 0.05, ** = p < 0.01; *** = p < 0.001).

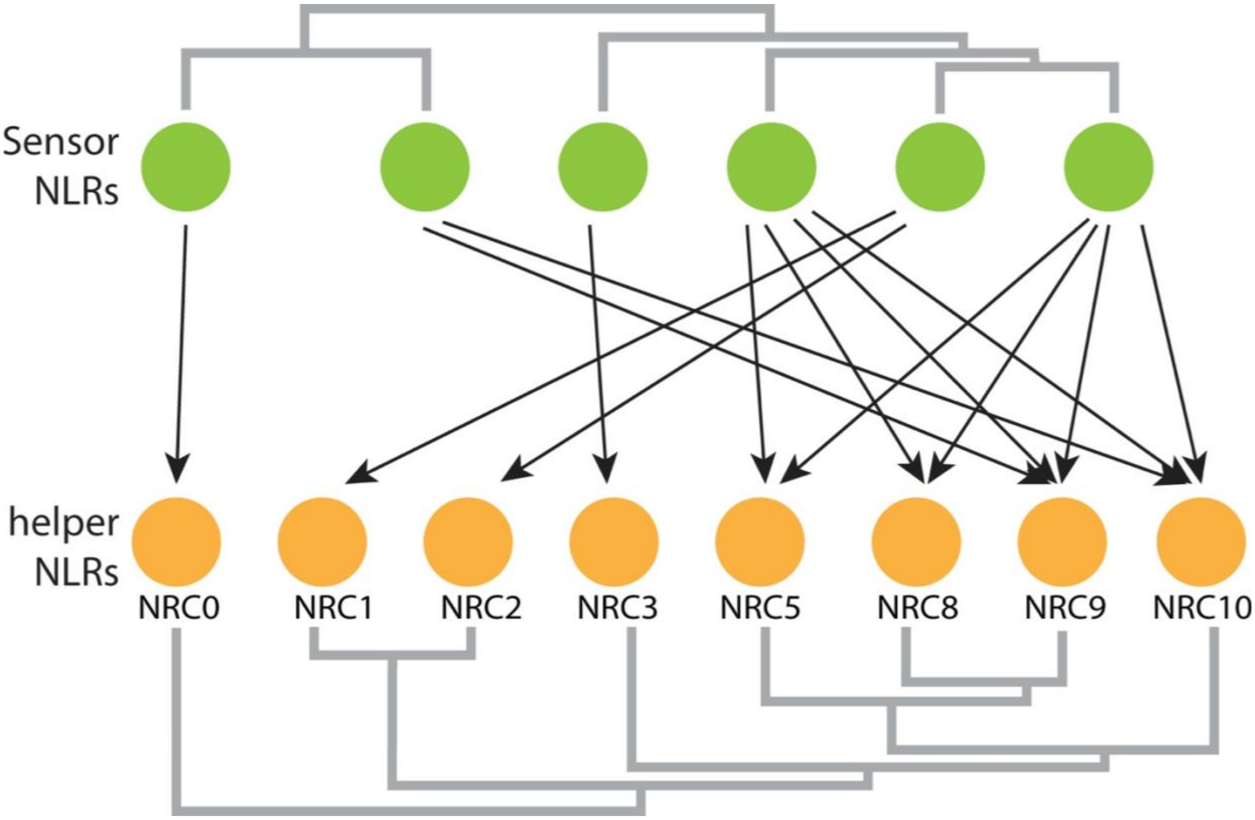


**Supplemental Fig. S8.** Water spinach possesses a complex NRC network. NRC0 can be specifically triggered by the NRC0-dependent sensor NLR to induce cell death. Certain sensor NLRs signal through Con-IaNRC1 or Con-IaNRC3 to induce cell death. Additionally, some sensor NLRs are capable of signalling through a few other NRC helpers to induce cell death.

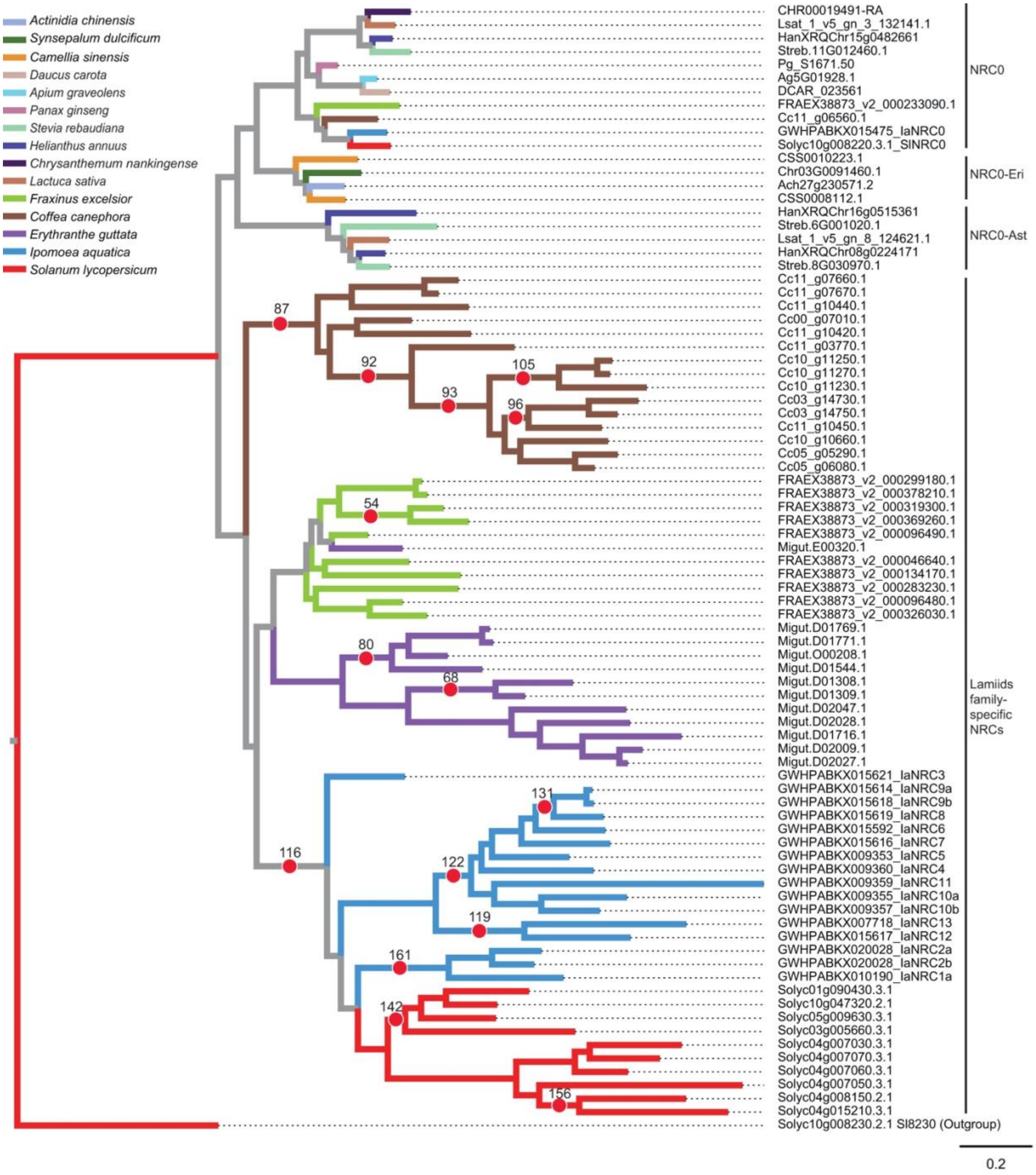


**Supplemental Fig. S9.** Several nodes in the phylogenetic tree of lamiids family-specific NRC subclades show diversifying selection. Full-length sequences of the NRC family from 15 selected asterids were used to generate the phylogenetic tree. The aBSREL analysis was used to detect internal branches with episodic diversifying selection, based on the Likelihood Ratio Test with a significance threshold set at $p \leq 0.05$. The scale bar indicates the evolutionary distance in amino acid substitution per site. The red dots indicate the 15 nodes showing diversifying selection and the numbers next to the red dots are the corresponding node numbers in the Supplemental Data Set 3.

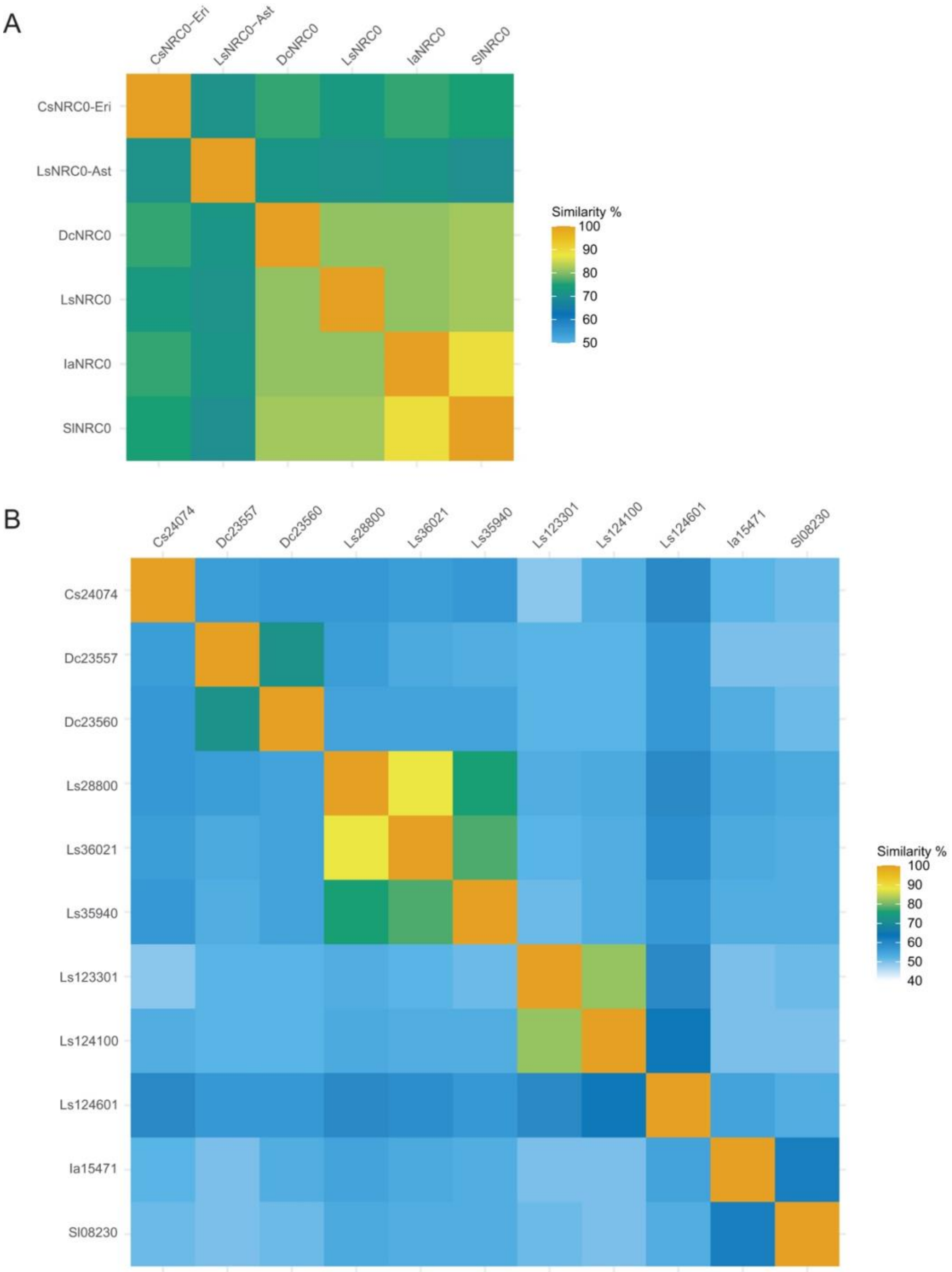


**Supplemental Fig. S10.** Pairwise sequence similarity comparisons of NRC0 subclades members and NRC-S tested in Figure 6. Pairwise sequence comparisons of (A) NRC0 subclades and (B) NRC0-dependent sensor NLRs were performed using BLASTP and the percentage of similarity (positive) was reported for each comparison.

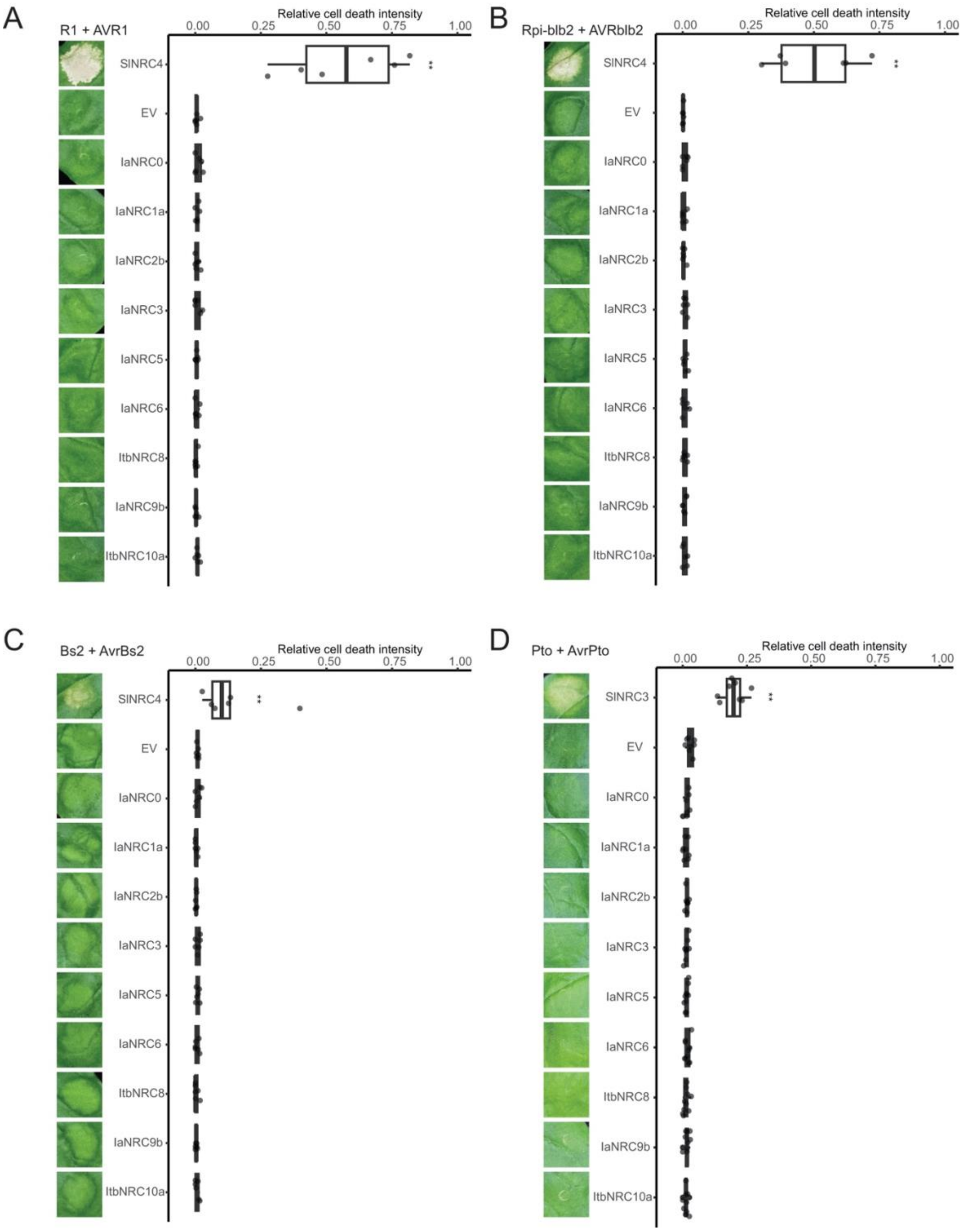


**Supplemental Fig. S11.** Solanaceous sensor NLRs R1, Rpi-blb2, Bs2, and Prf can not signal through any of the *Ipomoea* NRC tested. Solanaceous sensor NLRs **A)** R1, **B)** Rpi-blb2, **C)** Bs2, and **D)** Prf (Pto) were co-expressed with the corresponding AVRs and Con-IaNRCs in *N. benthamiana* leaves. Cell death phenotypes were recorded at 5 dpi. The dot plot represents the cell death quantification analysed by UVP ChemStudio PLUS. The bold line in the boxplots represents the medium, the box edges represent the 25th and 75th percentiles, and the whiskers extend to the most extreme data points no more than 1.5x of the interquartile range. Statistical differences were examined by paired Wilcoxon signed rank test (* = p < 0.05, ** = p < 0.01).

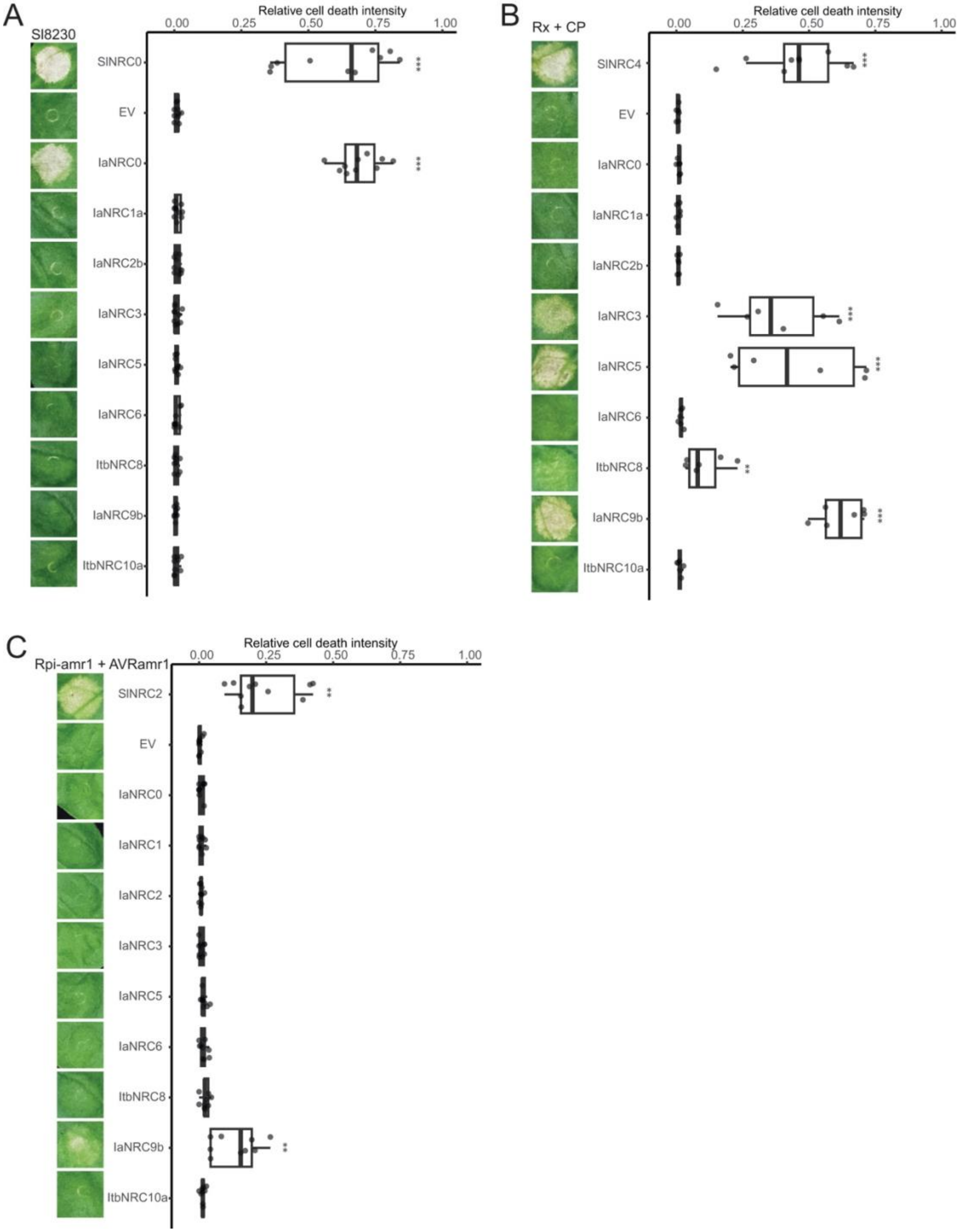


**Supplemental Fig. S12.** Solanaceous sensor NLR Rx, Rx2 and Rpi-amr1 can signal through some NRCs from *Ipomoea*. Solanaceous sensor NLR **A)** Sl8230 (Solyc10g008230), **B)** Rpi-amr1, **C)** Rx, and were made into autoactive or co-expressed with the corresponding AVRs and Con-IaNRCs in *N. benthamiana* leaves. Cell death phenotypes were recorded at 5 dpi. The dot plot represents the cell death quantification analysed by UVP ChemStudio PLUS. The bold line in the boxplots represents the medium, the box edges represent the 25th and 75th percentiles, and the whiskers extend to the most extreme data points no more than 1.5x of the interquartile range. Statistical differences were examined by paired Wilcoxon signed rank test (* = $p < 0.05$, ** = $p < 0.01$; *** = $p < 0.001$).

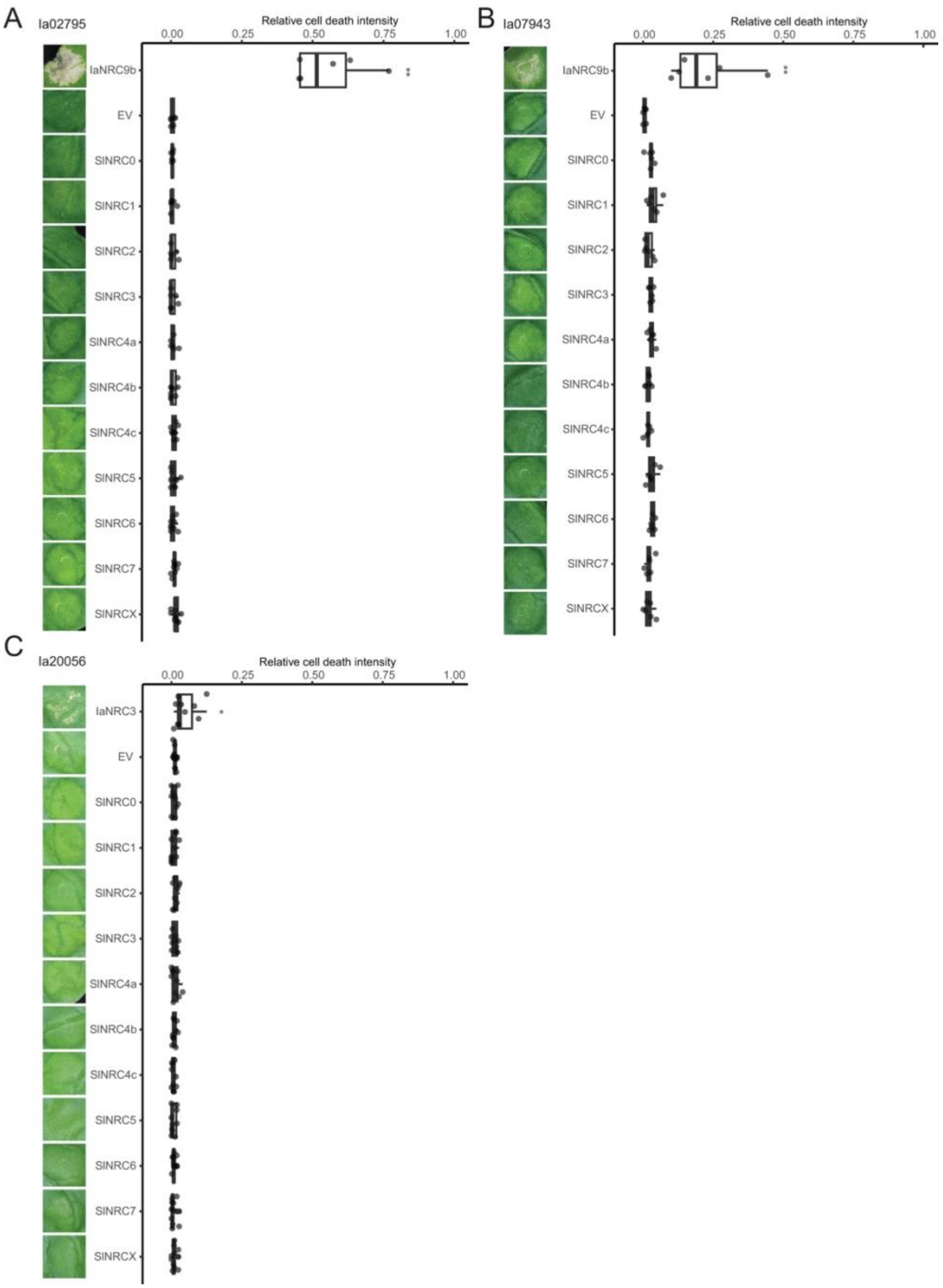


**Supplemental Fig. S13.** Some *I. aquatica* sensor NLRs can not signal through any of the tomato NRCs tested. Sensor NLR from *I. aquatica* **A)** Ia02795, **B)** Ia07943, and **C)** Ia20056 were made into autoactive (D to V mutation in the MHD motif) and then co-expressed with Sol-SlNRCs in *N. benthamiana.* Cell death phenotypes were recorded at 5 dpi. The dot plot represents the cell death quantification analysed by UVP ChemStudio PLUS. The bold line in the boxplots represents the medium, the box edges represent the 25th and 75th percentiles, and the whiskers extend to the most extreme data points no more than 1.5x of the interquartile range. Statistical differences were examined by paired Wilcoxon signed rank test (* = p < 0.05, ** = p < 0.01; *** = p < 0.001).

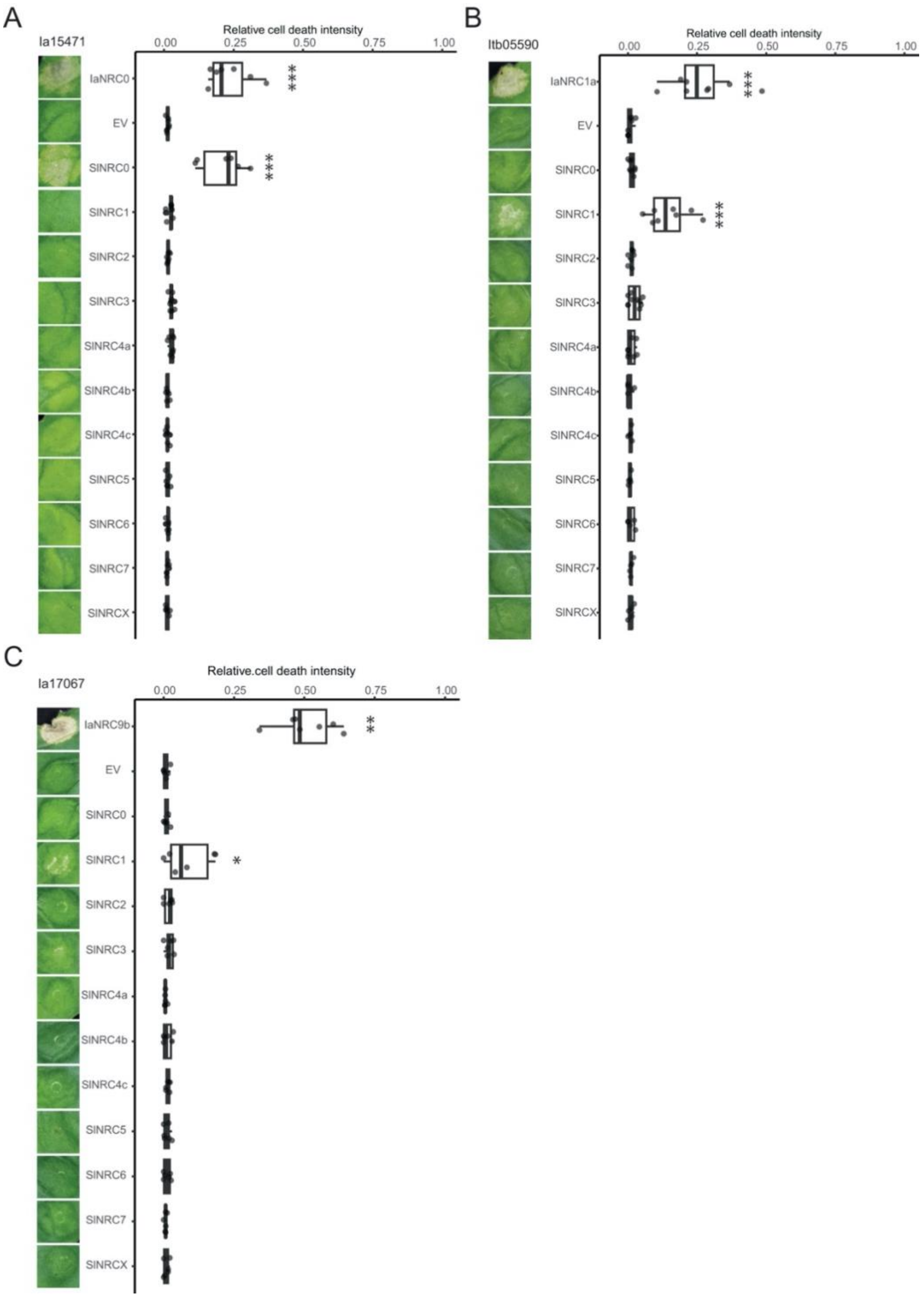


**Supplemental Fig. S14.** Some *Ipomoea* sensor NLR can signal through NRC1 of tomato. Sensor NLR from *Ipomoea* **A)** Ia15471, **B)** Itb05590, and **C)** Ia17067 were made into autoactive (D to V mutation in the MHD motif) and then co-expressed with Sol-SlNRCs in *N. benthamiana.* Cell death phenotypes were recorded at 5 dpi. The dot plot represents the cell death quantification analysed by UVP ChemStudio PLUS. The bold line in the boxplots represents the medium, the box edges represent the 25th and 75th percentiles, and the whiskers extend to the most extreme data points no more than 1.5x of the interquartile range. Statistical differences were examined by paired Wilcoxon signed rank test (* = p < 0.05, ** = p < 0.01; *** = p < 0.001).

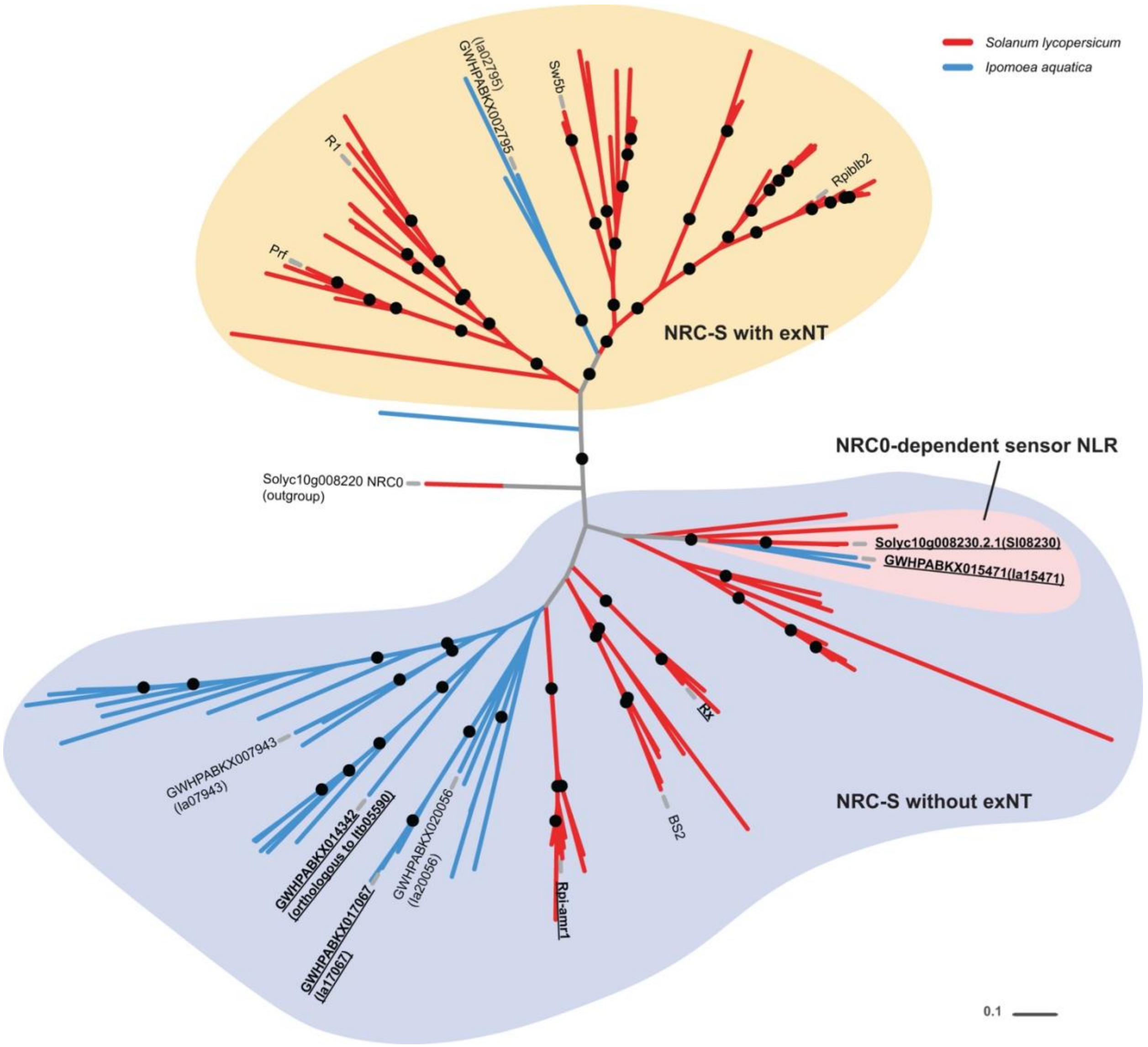


**Supplemental Fig. S15.** Phylogenetic analysis of NRC-S from tomato and *I. aquatica***.** Major nodes with bootstrap values over 70 are indicated with black dots. The scale bar indicates the evolutionary distance in amino acid substitution per site. NRC-S with or without N-terminal extension (exNT) and NRC0-dependent sensor NLRs are highlighted with different colour backgrounds. NLRs showing cross-compatibility with NRC-H from the other species are underlined.
